# The spatial structure of the tumor immune microenvironment can explain and predict patient response in high-grade serous carcinoma

**DOI:** 10.1101/2024.01.26.577350

**Authors:** Lucy Van Kleunen, Mansooreh Ahmadian, Miriam D Post, Rebecca J Wolsky, Christian Rickert, Kimberly Jordan, Junxiao Hu, Jennifer K. Richer, Nicole A. Marjon, Kian Behbakht, Matthew J. Sikora, Benjamin G. Bitler, Aaron Clauset

**Affiliations:** Department of Computer Science, University of Colorado, Boulder, USA; Department of Biostatistics and Informatics, Colorado School of Public Health, University of Colorado Anschutz Medical Campus, Aurora, CO, USA; Department of Pathology, The University of Colorado Anschutz Medical Campus; Department of Immunology and Microbiology, The University of Colorado Anschutz Medical Campus; Department of Pediatrics, Cancer Center Biostatistics Core, University of Colorado Anschutz Medical Campus, CO, USA; Department of OB/GYN, The University of Colorado Anschutz Medical Campus; BioFrontiers Institute, University of Colorado, Boulder, CO, USA; Santa Fe Institute, Santa Fe, NM, USA

## Abstract

Despite ovarian cancer being the deadliest gynecological malignancy, there has been little change to therapeutic options and mortality rates over the last three decades. Recent studies indicate that the composition of the tumor immune microenvironment (TIME) influences patient outcomes but are limited by a lack of spatial understanding. We performed multiplexed ion beam imaging (MIBI) on 83 human high-grade serous carcinoma tumors — one of the largest protein-based, spatially-intact, single-cell resolution tumor datasets assembled — and used statistical and machine learning approaches to connect features of the TIME spatial organization to patient outcomes. Along with traditional clinical/immunohistochemical attributes and indicators of TIME composition, we found that several features of TIME spatial organization had significant univariate correlations and/or high relative importance in high-dimensional predictive models. The top performing predictive model for patient progression-free survival (PFS) used a combination of TIME composition and spatial features. Results demonstrate the importance of spatial structure in understanding how the TIME contributes to treatment outcomes. Furthermore, the present study provides a generalizable roadmap for spatial analyses of the TIME in ovarian cancer research.

## INTRODUCTION

High grade serous carcinoma (HGSC) of the ovary, fallopian tube, and peritoneum is the gynecologic malignancy with the highest mortality rate (1,2). Over the last three decades there has been little improvement in the survival rate for patients diagnosed with HGSC, due in part to limited therapeutic options beyond chemotherapy, poor early detection rates, and a limited understanding of both the pathogenesis and the role of the tumor microenvironment. To further understand the drivers of HGSC and therapy response, several studies have examined patients who are disease-free 10 years after initial treatment (3). Long-term survival has been partially attributed to an enhanced anti-tumor immune response (4,5), indicating a clinical need to further define the tumor immune microenvironment (TIME) and elucidate its influence on patient outcomes.

Although HGSC often has a high degree of immune infiltrates, including macrophages that can compose up to 50% of all immune cells in the TIME (6), immune therapies have had limited impact on improving outcomes for individuals with HGSC (7). Prior studies of the HGSC TIME have discovered that selective immune cell infiltration often correlates with improved patient outcomes. Specifically, infiltration of CD3+ T cells and CD19+ B cells is associated with an average 62-month and 6-month survival benefit, respectively (8,9). In contrast, an increased density of CD163+ tumor associated macrophages within the TIME correlates with worse progression free survival (PFS) (10). Recently, spatial transcriptomics have proven to be a powerful tool to characterize the architecture of HGSC tumors, but these studies are currently performed with a limited spatial resolution (i.e., not single cell). These studies are also limited by their dependence on RNA expression (11–13). On the other hand, single cell sequencing of HGSC tumors provides significantly improved resolution of the TIME but is limited by the lack of associated spatial context (14). Recent studies have demonstrated that, beyond TIME composition, the spatial organization of the TIME, including the proximity of macrophages, B cells, and CD4+ T-cells to tumor cells significantly correlates with survival outcomes (15). However, these studies relied on a limited number of proteins to characterize the TIME spatial organization and thus were lacking simultaneous cell type identification, and the associations were not validated with modern large predictive models. Research in other types of cancer, such as melanoma, has shown that spatial features derived from single-cell image data are associated with treatment response (16).

In this study, we determined the prognostic power of the TIME’s spatial organization in explaining and predicting patient outcomes. Towards this end, we examined formalin-fixed paraffin-embedded (FFPE) tissue samples from 83 HGSC tumors from patients diagnosed with high grade serous carcinoma of the ovary, fallopian tube, and peritoneum with known outcomes with a multiplexed ion beam imaging (MIBI) system (17) and identified over 160,000 cells across 23 cell types. The resulting data set is one of the largest protein-based spatially intact, single cell analysis of any tumor type. Using survival and recurrence outcomes as an endpoint for 77 (69 primary and 8 recurrent) of the samples that met the inclusion criteria to produce spatial features, we performed modeling of 6 known clinical/immunohistochemical features (e.g., BRCA-status), 24 TIME composition features, 69 TIME spatial features, and 117 TIME (spatial) network features to assess their correlation with and relative importance for predicting patient outcomes. We found significant univariate correlations and high relative importance in high-dimensional predictive models for several features encoding TIME spatial organization. While we were unable to reliably predict out-of-sample overall survival (OS) outcomes with these features, we consistently predicted out-of-sample PFS, with the best model on average using a combination of features of the TIME composition and spatial organization. We demonstrate how moving beyond TIME composition to encode and assess features of TIME spatial organization, combined with a modern machine learning approach, can be used to improve hypothesis generation and testing to identify clinically relevant parameters for improving HGSC patient care.

## RESULTS

### Multiplexed imaging, cell segmentation, and phenotyping

We performed multiplexed imaging using a custom MIBI-TOF instrument (17) to produce a total of 83 images identifying 26 proteins (File D1), which were processed using Ionpath’s MIBI/O software and corrected (Table S1) and denoised (File D2). Multiplexed imaging data were preprocessed to remove noise and artifacts as described previously (26) prior to single-cell segmentation. In this preprocessing step, we used supervised pixel classification to generate a feature representation map for each image (Fig. 1A). We then applied a widely used pre-trained model (27) to perform whole-cell segmentation. This process identified about 160,000 cells with each FOV containing an average of ∼1934 single cells (s.d=556). The unsupervised clustering algorithm FlowSOM (28) was then employed, identifying 23 unique cell clusters (Fig. 1B,C, Fig. 2A). The cell type identity of each cluster was determined by comparing relative phenotypic marker signal intensities across clusters.

**Fig 1.**
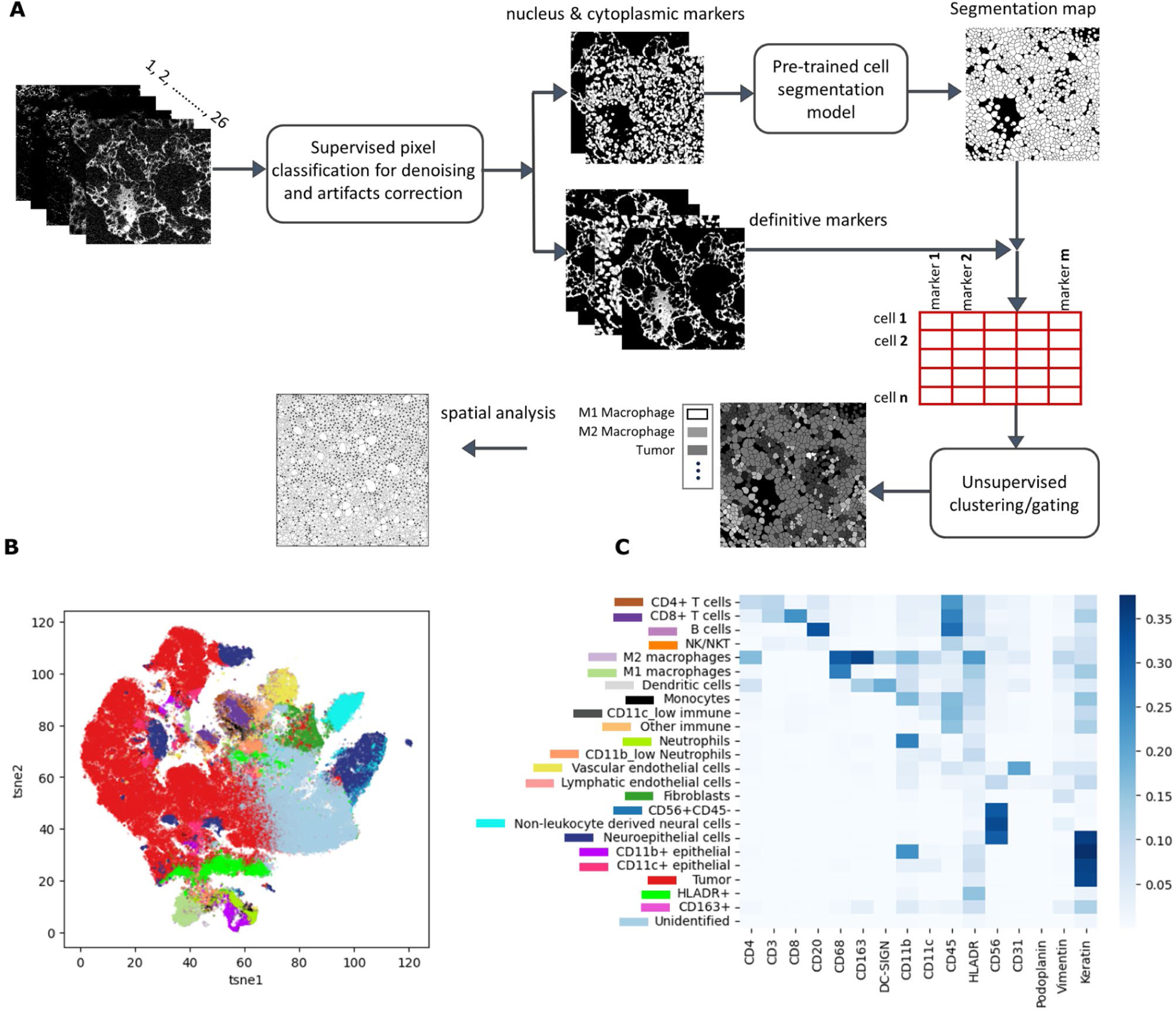
Cell segmentation and phenotyping. (A) Computational pipeline used for single-cell segmentation and cellular phenotyping of the MIBI imaging data. The process starts with pixel classification, where a pixel classifier distinguishes between two classes: Class I for desired signals and Class II for noise and artifacts. The classifier’s output produces feature representation maps with pixel values scaled from 0 to 1. A pretrained single-cell segmentation model is used for cell segmentation. Subsequently, marker expression within cell boundaries is quantified using the Class I feature representation maps. This data is organized into a single-cell information table, with cells listed in rows and marker expression levels in columns. Finally, unsupervised clustering algorithms utilize this single-cell information data to identify distinct cell types. (B) tSNE representation of the marker expression data of about 160k cells from the ovarian cancer tissue of 83 patients. Cell types were identified by clustering (represented in different colors). (C) Average marker expression per cluster is shown for the identified cell types, with colors indicating their corresponding cluster in the *tSNE* representation.

**Fig 2.**
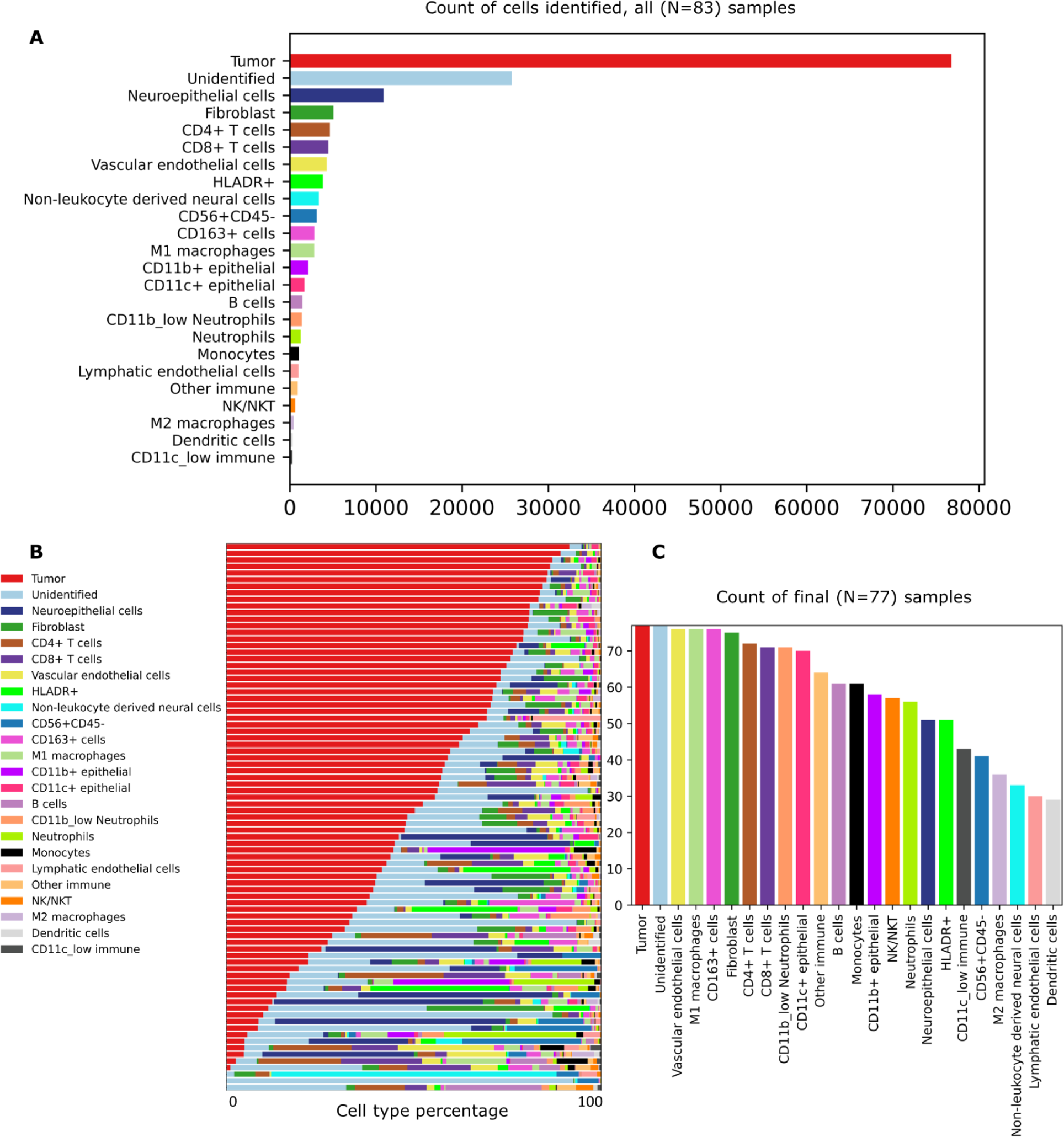
TIME composition across samples. (A) Cell type counts across all 83 samples. (B) Cell type percentages summarized across the 83 original samples, sorted by decreasing tumor cell percentage. (C) Counts of the final 77 samples included in the analysis in which each of the cell types were found.

### Generating TIME composition features

We first examined the TIME composition of the samples in terms of the relative frequency of cell types. This composition spanned 24 features of the samples exclusive of spatial organization, comprising 23 cell types and the population of unidentified cells. We observed substantial variation in cell type frequencies across samples (Fig.2B). Tumor cells were the most prevalent cell type, representing on average 47.8% of the cells in each sample (range 0% to 91.6%). The next most common cell types on average were neuroepithelial cells (mean 6.4%, range 0% to 61.5%; vs. tumor cell percentage, Pearson correlation coefficient *r*=-0.328, false discovery rate adjusted two-sided *p*=0.012). All other cell types varied from 0% to 3.3% of the cells on average, though these percentages could vary dramatically between samples, often in relation to tumor cell percentage. Other cell types with false discovery rate adjusted significant correlation coefficients with tumor cell percentages were CD8+ T cells (*r*=-0.311, *p*=0.013), CD4+ T cells (*r*=-0.379, *p*<0.001), NK/NKT cells (*r*=-0.303, *p*=0.014), CD56+CD45-cells (*r*=-0.401, *p*<0.001), vascular endothelial cells (*r*=-0.29, *p*=0.018), B cells (*r*=-0.339, *p*=0.009), monocytes (*r*=-0.288, *p*=0.018), CD11c^low^ immune cells (*r*=-0.28, *p*=0.021), neutrophils (*r*=-0.257, *p*=0.036), and CD11c+ epithelial cells (*r*=0.328, *p*=0.009). All other cell types did not have significant correlations (File D3). Some cell types such as dendritic cells (DC) and CD11c^low^ immune cells were always rare, if present in a sample.

We excluded some samples from further analysis based on cell type percentages and two exclusion criteria (Fig. S1). Unidentified cells represented on average 16.7% of the cells in each sample (range 0.5% to 92.6%; *r*=-0.498, *p*<0.001). Samples 26 and 45 were excluded because they were outliers with unidentified cell percentages over 65%. Samples 27 and 29 were excluded because they had no identified tumor cells (sample 45 also met this exclusion criteria). We determined that samples with a high percentage of unidentified cells or no identified tumor cells were unable to produce spatial features about the interactions between cells of different types, and in particular interactions with tumor cells. In the two cases in which there were two samples from the same patient, we chose to keep the sample with a lower unidentified cell percentage in the final analysis, thus excluding samples 19 and 35. This choice ensured that our final dataset included at most one sample from each patient in the analysis linking generated features to patient outcomes. In total, we excluded 6 samples from the final analysis, leading to a final dataset of 77 samples.

Most cell types were not represented across all images in the final dataset (Fig. 2C). Tumor cells were identified in every sample, and vascular endothelial cells, M1 macrophages, CD163+ cells, and Fibroblast cells were identified in almost every sample. Some cell types were rarer, particularly M2 macrophages, non-leukocyte derived neural cells, lymphatic endothelial cells, and dendritic cells were identified in fewer than half of the samples.

### Generating spatial features of the TIME based on nearest neighbor distances

For each sample in the final dataset, we generated a set of 69 features that characterize each sample’s spatial structure, following the approach from Moldoveanu et al. (2022) (16). First, we generated the median Euclidean distance from three distinct cell types (“focal cell types”) that have been reported to be important in the HGSC TIME (tumor cells, M1 macrophages, and vascular endothelial cells) in each sample to their nearest neighbors of each other cell type. While there have been few studies interrogating the spatial features of the TIME, previous work indicates that the spatial proximity between cell types correlates with HGSC survival outcomes (15). Previous results on composition (10,38,39), led us to focus on M1 macrophages and vascular endothelial cells as focal cell types for generating spatial and network features along with tumor cells in our study. Vascular endothelial cells and M1 macrophages were also both detected in nearly all (98%, only missing in one sample each respectively) samples.

Tumor cells, vascular endothelial cells, M1 macrophages, CD163+ cells, and fibroblasts, which were some of the most common cell types across samples, were closer (average median nearest neighbor distance under 90 μm) than other cell types to all three focal cell types (Fig.3A-C). In comparison, the cell types that were in fewer samples (e.g., M2 macrophages, non-leukocyte derived neural cells, lymphatic endothelial cells, dendritic cells) were found on average further away from the three focal cell types. B cells had the highest mean nearest neighbor distance across samples to all three focal cell types (197.2 μm to Tumor cells, 190.4 μm to M1 macrophages, and 174.2 μm to vascular endothelial cells, respectively).

**Fig 3.**
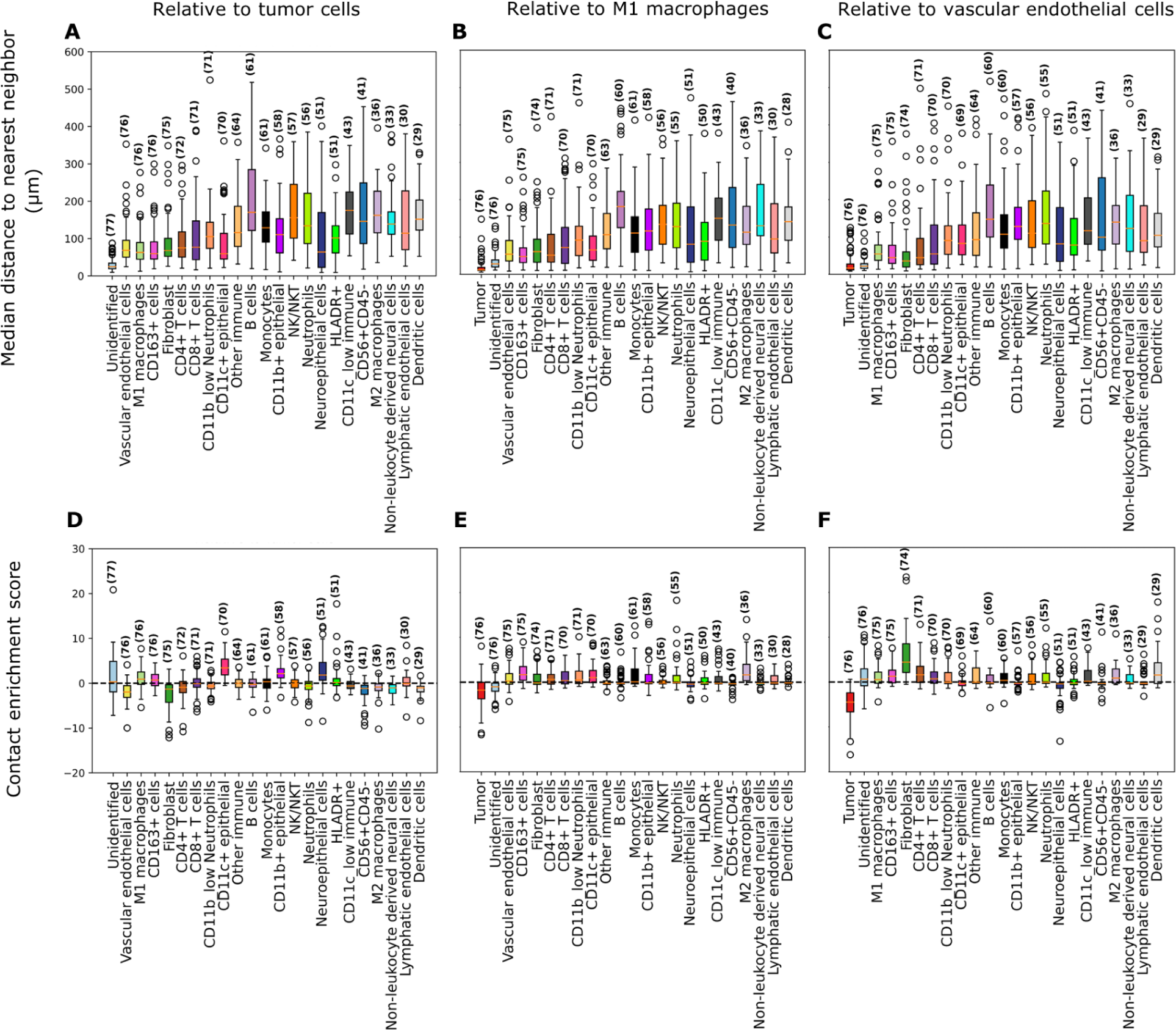
Median nearest neighbor distance (spatial) and contact enrichment (network) features relative to three focal cell types. (A-C) Median nearest neighbor distance for each other cell type to tumor cells, M1 macrophages, or vascular endothelial cells (μm). (D-F) Contact enrichment scores relative to tumor cells, M1 macrophages, or vascular endothelial cells for each of the other cell types. Positive scores indicate more contacts than expected at random, 0 the same number, and negative scores fewer contacts than expected at random. All bar plots show features aggregated across samples in which the relevant cell type is found. Cell types are indicated on the x-axis and the number of samples in which this cell type is found is shown in parentheses. Samples are excluded from the features calculated relative to M1 macrophages and vascular endothelial cells respectively when samples are missing the respective focal cel l type. In all subplots cell types are ordered based on how commonly they were found across samples in descending order.

### Generating features of the TIME based on spatial network representations

We next created spatial network representations of the samples by connecting spatially proximate cells using Delaunay triangulation and then trimming edges that were above a threshold of 50 pixels (∼24.4 μm) (Fig. S2A-C). Cells were thus found 15-50 pixels (∼7.3-24.4 μm) away from their spatial neighbors in the networks, with some variation in the median distances between spatial network neighbors based on their cell type (Fig. S3). Using these spatial network representations, we generated 117 TIME network features for each sample. The first subset of these features represented the mean size of connected regions of each cell type in each sample (Fig. S2D). These spatially connected regions in the TIME may indicate the existence of spatially extended structures of a single cell type (which may indicate the level of tumor infiltration, per Keren et al. 2018 (30)). Tumor cells were most often found in large, connected regions with 50% located in regions of 1071 cells or greater. Neuroepithelial cells were also found in relatively large, connected regions, with 50% found in regions of 226 cells or greater. Cells of all other types were most often found in relatively small regions ranging from 1 to 14 cells (Fig. S4A). Most regions of any cell type were only 1 or 2 cells large. The largest maximum region sizes were for tumor cells (2190 cells), Neuroepithelial cells (1288 cells), CD11b+ epithelial cells (349 cells), and CD4+ T cells (202 cells), while all other cell types had maximum region sizes under 200 cells (Fig. S4B).

We used the spatial network representations to compute contact enrichment scores, following prior work (30) (16) (31) to quantify the extent to which network-neighbors of focal cell types might differ from what should be expected at random. These scores capture similar proximity information as the median nearest neighbor distance features, but control for the proportion of cells of each type by keeping these proportions fixed during computation. Moreover, these scores quantify direct interactions between cell types.

Vascular endothelial cells, fibroblasts, and CD56+CD45-cells had fewer contacts with tumor cells than expected based on random sampling (null expectations), whereas CD11c+ epithelial cells, CD11b+ epithelial cells, and neuroepithelial cells often had more contacts than expected. Most other cell types varied across samples with many contact enrichment scores close to 0, and thus matching null expectations (Fig. 3D). Most of the cell types showed slightly more contacts with M1 macrophages than expected at random (contact enrichment scores > 0), with the exception of tumor cells and neuroepithelial cells, which tended to have fewer (contact enrichment scores < 0) and other immune cells, B cells, NK/NKT cells, CD56+CD45-cells, non-leukocyte derived neural cells, and dendritic cells which tended to have contact enrichment scores with M1 macrophages close to 0 (Fig. 3E). Contact enrichment scores with vascular endothelial cells were also slightly positive for most cell types and negative for tumor cells. Fibroblasts had more contacts with vascular endothelial cells than expected at random and cell types with slightly negative or varying vascular endothelial contact enrichment scores included CD11c+ epithelial cells, neuroepithelial cells, CD56+CD45-cells, and lymphatic endothelial cells (Fig.3F).

Finally, we generated assortativity coefficients from the spatial networks, which measure the extent to which cells tend to be network neighbors with cells of the same type as opposed to neighbors of any other type. These features capture similar information about large-scale structure and tumor infiltration as the mean region size, but better account for random variation. Tumor cells had the highest mean assortativity coefficient (0.37). We did not observe any cells exhibiting a negative assortativity coefficient (disassortative mixing), in which cells of a given cell type would be less likely to be network neighbors with same-type cells and more likely to be neighbors of different-type cells. We did, however, see large variation across samples in the magnitude of the assortativity coefficient for many cell types. For instance, the tumor cells displayed a large range of assortativity coefficients, which may indicate that the tumors in some samples were more infiltrated by other cells (Fig. 4).

**Fig 4.**
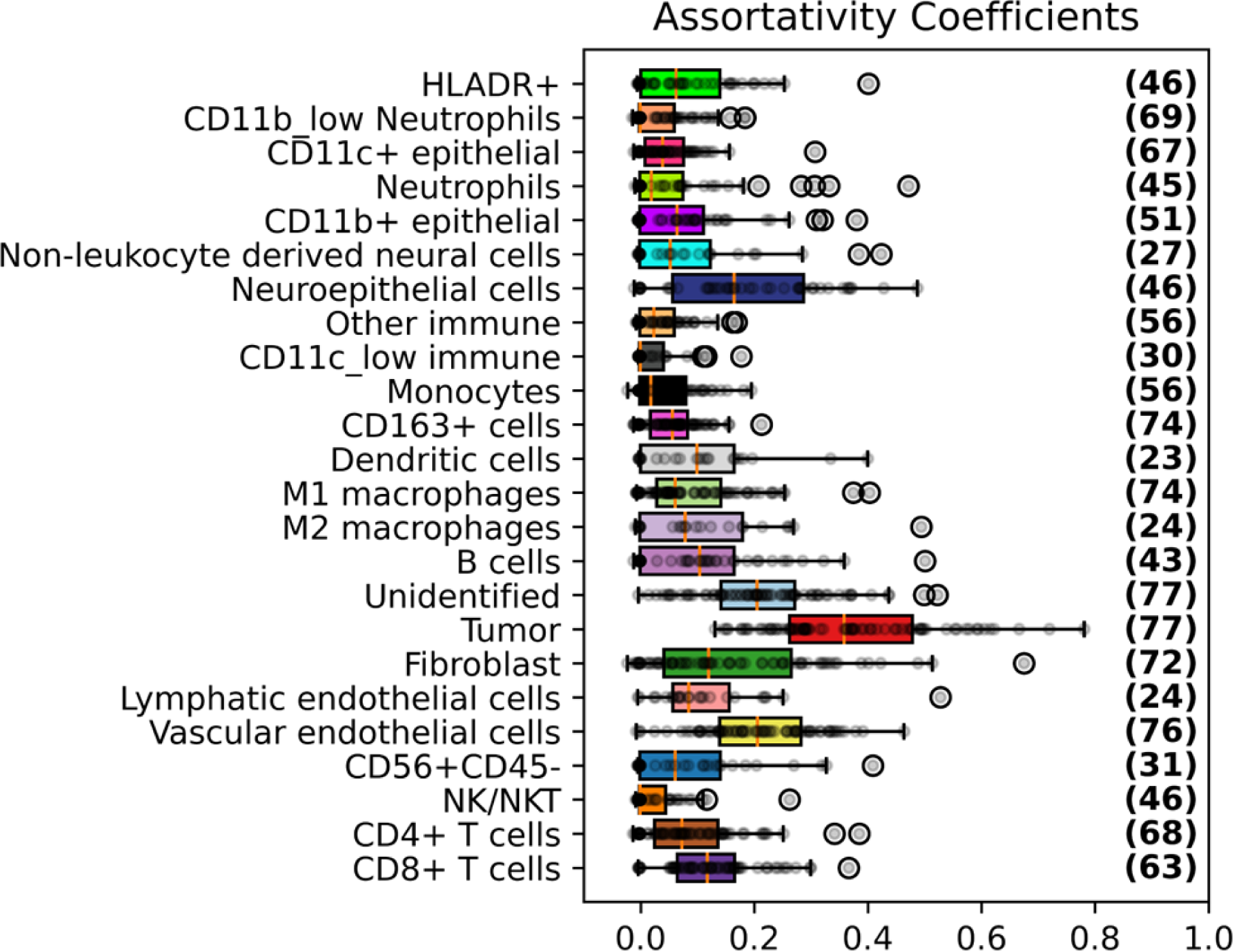
Assortativity coefficient (network) features. Assortativity coefficients for each cell type indicating their tendency to cluster with cells of the same type rather than cells of a different type, aggregated across samples including that cell type (the number of which is indicated in parentheses).

### Linking in-sample patient outcomes to TIME features in univariate and multivariate Cox regressions

We next explored the relationship between generated features of the samples and two time-to-event outcomes: overall survival (OS) and progression-free survival (PFS) (Fig. S5, Fig. S6). We define OS as the time from initial diagnosis based on tissue biopsy and imaging or a serum biomarker (CA125) to death. Patient data without observation of death are censored at the last known patient visit. PFS is defined as the time from initial diagnosis to first known disease recurrence. Patient data without observation of disease recurrence are censored at death or the last known patient visit, whichever occurs first.

We performed Univariate Cox regressions for all the generated TIME composition, spatial, and network features as well as 6 additional clinical/immunohistochemical features: age, BRCA mutational status, H3K14Ace status, ATF6 status, DUSP1 status, and CBX2 status (see Fig. S7 for clinical/immunohistochemical feature distributions). All covariates except BRCA mutational status and age were first normalized (z-score scaled) so that they had a mean of 0 and a standard deviation of 1. Results limiting the dataset to only primary tumor samples can be found in Figure S8 and File D4.

For OS, we found significant univariate results (p<0.05) for 25 features associated with worse prognosis and 3 features associated with better prognosis (Fig. 5A, see File D5 for full results). For PFS, we found significant results (p<0.05) for eight features associated with worse prognosis and three features associated with better prognosis (Fig. 5B, see File D5 for full results). None of the clinical/immunohistochemical attributes were significant for either outcome variable.

**Fig 5.**
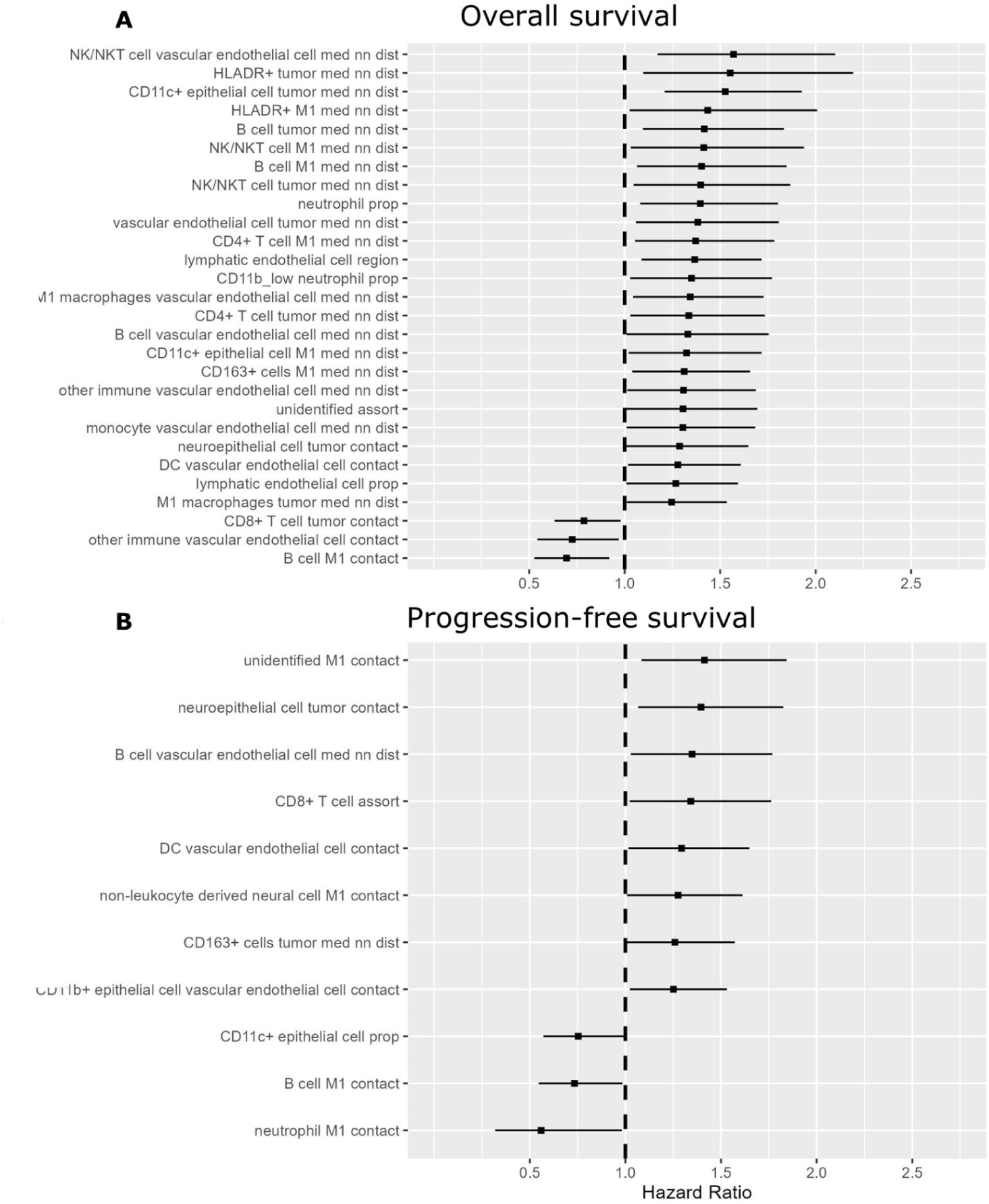
Univariate Cox regression results. Covariates found to be significant in Univariate Cox regressions for (A) OS and (B) PFS outcomes. Covariates are listed in descending order by hazard ratio. Hazard ratios are displayed with 95% confidence intervals, and a hazard ratio of 1 is indicated with a dashed line.

Of the significant features, a majority were related to proximity between cells of particular types – median nearest neighbor distance features accounted for 18 of the 28 significant features for OS and two of the 11 significant features for PFS. Contact enrichment features accounted for five of the 28 significant features for OS and seven of the 11 significant features for PFS. Relatively fewer of the significant features related to composition or the tendency for cells of the same type to cluster together – three composition features were significant for OS and one composition feature was significant for PFS, one mean region size feature was significant for OS, and none were significant for PFS, and one assortativity coefficient feature was significant for each of OS and PFS.

As a robustness check, we trained a set of reduced models, in the form of multivariate Cox regressions on only the top five features for each outcome variable ranked by *p*-value, with and without adjusting for the 6 clinical/immunohistochemical features (Table S2, S3). In the adjusted multivariate model for OS, none of the five top features remained significant, while in the reduced model the only feature that remained significant was the median NK/NKT cell nearest neighbor distance to vascular endothelial cells (Hazard Ratio [HR] =1.66, p=0.009). For PFS, the only feature that remained significant in the adjusted model was the contact enrichment score between unidentified cells and M1 macrophages (HR=1.63, p=0.010), while in the reduced model four of the top five features remained significant while the contact enrichment score between unidentified cells and M1 macrophages was not significant (HR=1.32, p=0.059).

### Predicting out-of-sample patient outcomes using TIME features in random forests

We next evaluated how spatial and/or network features of the tumor microenvironment could be used together with clinical/immunohistochemical attributes and TIME composition features to predict patient outcomes out-of-sample. We split both OS and PFS outcome variables at their respective medians to consider a simple binary classification task of low or high OS or PFS. We grouped features into 4 categories according to their respective processes of derivation: (i) clinical/immunohistochemical, (ii) composition, (iii) spatial, and (iv) network features. We considered all 15 possible combinations of the four feature categories to evaluate what combination of information produced the best out-of-sample predictive performance (Fig. 6A).

**Fig 6.**
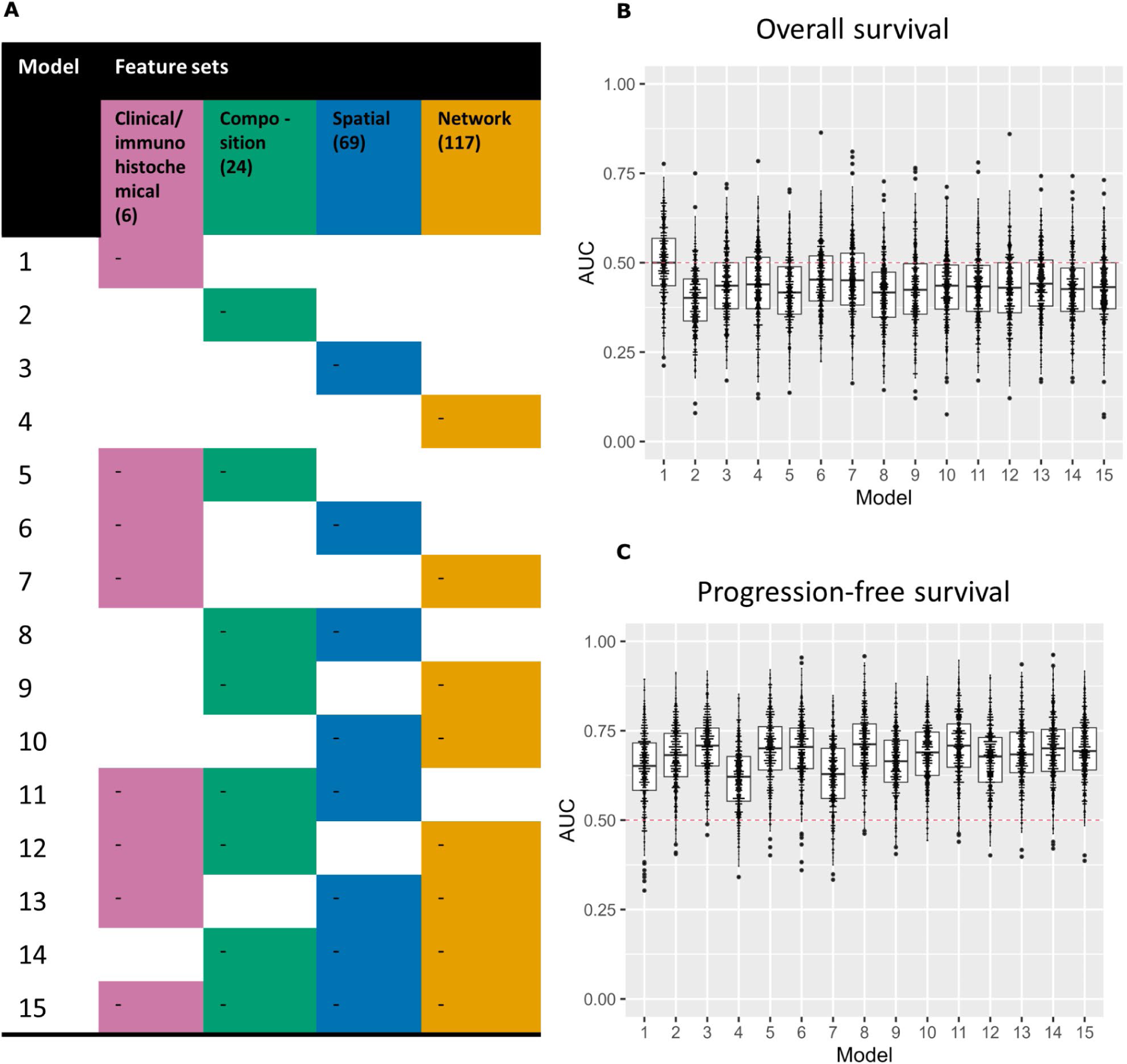
Random forest predictive performance results. (A) 15 models were trained and evaluated with different combinations of four feature categories, as shown here. Predictive performance results, based on the AUC statistic are displayed for the 15 models summarized across 500 iterations of training and evaluation for (B) OS and (C) PFS outcomes. A red dashed line is displayed at an AUC value of 0.5, which represents the cut-off above which the model performs better than a random guess.

For each model, we repeatedly (N=500) trained a random forest model on a training set of 70% of the samples, randomly sampled while balancing outcome labels, and evaluated each model’s predictive accuracy by using the remaining 30% as a test set. We report average out-of-sample predictive performance results, based on the AUC (Area Under the Receiver Operating Characteristics curve) statistic (36) across these 500 evaluations. A value of AUC=0.5 indicates a classification that performs no better or worse than a random guess, while an AUC=1 indicates perfect performance. Across these predictive models, we found better-than-random performance on average, with AUC>0.5 for PFS but not for OS (Fig. 6B, C, Table S4). All models for PFS achieved mean AUC values over 0.6. The model that best predicted PFS was model eight, with AUC 0.711 ± 0.10 (mean ± stddev) based on combining composition and spatial features. This performance was followed closely by model 11 (0.707 ± 0.09) and model 3 (0.703 ± 0.08), which used only spatial and a combination of clinical/immunohistochemical, composition, and spatial features, respectively. Models containing network features performed slightly worse (models 4, 7, 9, 10, 12, 13, 14, 15; average AUC= 0.668±0.03) than models with clinical/immunohistochemical features (models 1, 5, 6, 7, 11, 12, 13, 15; average AUC=0.678±0.03), composition features (models 2, 5, 8, 9, 11, 12, 14, 15, average AUC=0.690), and spatial features (models 3, 6, 8, 10, 11, 13, 14, 15; average AUC=0.698±0.01). The model containing all features (model 15) achieved an AUC of 0.697 ± 0.09. All the models predicting OS achieved mean AUC<0.5, indicating that on average the models did not outperform a random guess, i.e., they predicted in the incorrect direction (Fig. 5B). Similar performance results were found for models trained only on primary tumor samples (n=69), although composition features were relatively more helpful in predicting PFS, such that the top three models were model five (0.729 ± 0.09), model two (0.719 ± 0.09), and model eight (0.714 ± 0.09) (Fig. S9, Table S5).

Using model 15, which was trained on all four feature categories, we generated hypotheses of which particular TIME composition, spatial, and network features were relatively more useful for predicting OS and PFS patient outcomes. We accomplished this goal by calculating and comparing the Gini importance scores (37) for each feature in model 15. We note that importance scores do not indicate the direction of a feature’s relationship with a patient’s outcome, and instead only indicate its relative utility in predicting the outcome value. We found evidence for a subset of features, spanning all four categories, that were relatively more important for predicting patient outcomes (Fig. 7A,B). Feature importance results limiting the dataset to only primary tumor samples can be found in Figure S10 and File D6.

**Fig 7.**
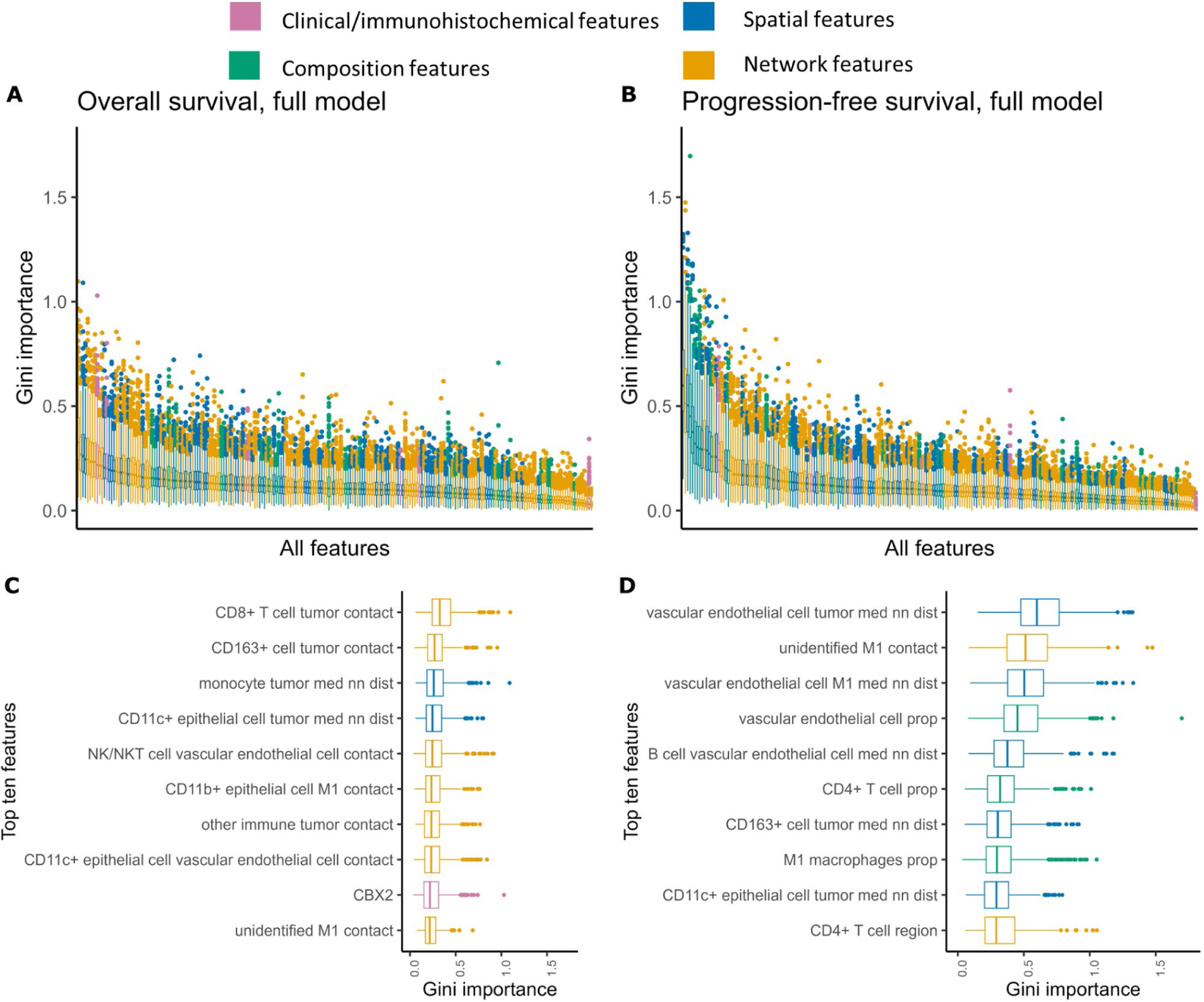
Aggregate feature importance results. Gini importance scores, aggregated across 500 random forest training runs for the model including all features, sorted by median importance score and colored by feature type for (A) OS and (B) PFS outcomes. (C-D) Top ten features by median importance score for each outcome across 500 random forest training runs, colored by feature type.

Ranking features by their median Gini importance score across the 500 evaluations, we found that the top ten features for predicting OS included seven contact enrichment network features, two spatial features, and one clinical/immunohistochemical feature: (i) the contact enrichment score between CD8+ T cells and tumor cells, (ii) the contact enrichment score between CD163+ cells and tumor cells, (iii) median monocyte cell nearest neighbor distance to tumor cells, (iv) median CD11c+ epithelial cell nearest neighbor distance to tumor cells, (v) the contact enrichment score between NK/NKT cells and vascular endothelial cells, (vi) the contact enrichment score between CD11b+ epithelial cells and M1 macrophages, (vii) the contact enrichment score between other immune cells and tumor cells, (viii) the contact enrichment score between CD11c+ epithelial cells and vascular endothelial cells, (ix) CBX2 status, and (x) the contact enrichment score between unidentified cells and M1 macrophages (Fig. 7C, File D7). The contact enrichment score between CD8+ T cells and tumor cells were distinguished by a higher median feature importance score.

We found that the top ten features for predicting PFS included one contact enrichment network feature, one mean region size network feature, five spatial features, and three composition features: (i) median vascular endothelial cell nearest neighbor distance to tumor cells, (ii) the contact enrichment score between unidentified cells and M1 macrophages, (iii) median vascular endothelial cell nearest neighbor distance to M1 macrophages, (iv) vascular endothelial cell proportion, (v) median B cell nearest neighbor distance to vascular endothelial cells, (vi) CD4+ T cell proportion, (vii) median CD163+ cell nearest neighbor distance to tumor cells, (viii) M1 macrophage proportion, (ix) median CD11c+ epithelial cell nearest neighbor distance to tumor cells, and (x) the CD4+ T cell mean region size (Fig. 7D, File D7). In particular, for PFS the median vascular endothelial cell nearest neighbor distance to tumor cells was consistently ranked as more important across the iterations.

In alignment with the in-sample Cox regression results, we found that most of the top ten most important features for predicting out-of-sample patient outcomes (nine for OS and six for PFS) were either median nearest neighbor distances or spatial network contact enrichment scores, and thus related to the spatial proximity between cell types, and in general, features related to spatial proximity were more important (Fig. S11, Fig. S12).

### Results related to T cell, macrophage, B cell, and vascular endothelial cell spatial organization

Previous work has indicated that the presence of intratumoral T cells and the presence of T cells in ascites have been shown to correlate with better patient prognosis (8,15,40–42). High CD4+ T cell macrophage interaction has also been shown to significantly correlate with better prognosis when adjusted for clinical/immunohistochemical covariates (15). In our results, patients with NK/NKT and CD4+ T cells closer to M1 macrophages and tumor cells and NK/NKT cells closer to vascular endothelial cells had better OS, and the contact enrichment score between CD8+ T cells and tumor cells was the most important feature for predicting OS. For PFS, the same features were not significantly correlated with prognosis, though NK/NKT cell M1 median nearest neighbor distance was significantly correlated with PFS for only primary tumor samples (HR=1.43, p=0.038). We also saw a significant correlation for CD8+ T cell assortativity (HR 1.34, p=0.034, not significant for only primary tumor samples), and the CD4+ T cell mean region size was chosen as an important feature for the random forest predicting PFS and was significantly correlated with PFS for only the primary tumor samples (HR 0.73, p=0.046), indicating that T cell clustering patterns might have been more important for predicting PFS than T cell spatial proximity features.

Macrophages compose up to 50% of all immune cells in the TIME and are a highly plastic cell type (6). As opposed to M2-like macrophages, M1-like macrophages are proposed to be anti-tumorigenic and aid the adaptive immune cells in mounting an immune response (43). The M1/M2 ratio of macrophages in the ovarian cancer TIME is prognostic for overall and progression-free survival (10,38,44). For macrophages, our significant results all were related to M1 macrophages rather than related to M2 macrophages. We found that higher median M1 macrophage nearest neighbor distance to tumor cells (HR 1.25, p=0.039) or vascular endothelial cells (HR=1.34, p=0.021, not significant for only primary tumor samples) were significantly correlated with worse OS. Median vascular endothelial cell nearest neighbor distance to M1 macrophages was also chosen as an important feature by the random forest for predicting PFS.

We found, in alignment with previous results (15) that a higher contact enrichment score between B cell and M1 macrophages was significantly correlated with both better OS (HR=0.696, p=0.011) and PFS (HR=0.73, p=0.039), and that a larger median B cell nearest neighbor distance to M1 macrophages was significantly correlated with worse OS (HR=1.40, p=0.016, not significant for only primary tumor samples). Unlike in Steinhart et al. 2021 (15) we differentiated between M1 and M2 macrophages, replicating this result for the former and thus adding further cell type specificity to these findings. These findings highlight that interaction between B cells and M1 macrophages may lead to a better antitumor immune response after chemotherapy treatment potentially through increased macrophage-mediated antigen presentation to the B cells. We generally observed that B cells being further from M1 macrophages, vascular endothelial cells, and tumor cells corresponded to worse outcomes - higher median B cell nearest neighbor distance to tumor cells (HR=1.42, p=0.008) or vascular endothelial cells (HR=1.33, p=0.042, not significant for only primary tumor samples) also significantly correlated with worse OS, and median B cell nearest neighbor distance to vascular endothelial cells was also significantly correlated with worse PFS (HR=1.35, p=0.030) and was chosen as an important feature for predicting PFS.

A higher density of microvessels in the TIME has also previously been correlated with worse progression-free survival (39), and anti-angiogenic therapies (e.g., anti-VEGF) are a standard of care for ovarian cancer. In our results, OS was significantly correlated (HR=1.23, p=0.073) with the median nearest neighbor distance between vascular endothelial cells and tumor cells, as in a higher median nearest neighbor distance between these cell types conveyed a worse OS. Median vascular endothelial cell nearest neighbor distance to tumor cells was also chosen as the most important feature for predicting PFS.

## DISCUSSION

The current study provides a roadmap for further hypothesis generation and evaluation in ovarian cancer research, opening a range of possible directions for future work investigating the mechanisms by which TIME spatial organization drives clinical and biological differences.

Our results reinforce the importance of considering the spatial structure of the TIME to understand and predict HGSC disease progression and outcomes. We show that features encoding the spatial and network organization of the TIME help predict patient outcomes, and we find that the best predictive model for PFS includes a combination of TIME composition and spatial features. For example, we found several results related to CD163+ cells, e.g., higher median CD163+ cell nearest neighbor distance to M1 macrophages correlated with worse OS (HR=1.31, p=0.022) and higher median CD163+ cell nearest neighbor distance to tumor cells correlated with worse PFS (HR=1.26, p=0.042) and was chosen as an important feature for predicting PFS. CD163 is a scavenger receptor, and its expression is largely restricted to myeloid-derived cells, specifically monocytes and macrophages – it is often upregulated in response to inflammation and is associated with tumor promoting macrophages (45). While CD163+ cells in the ovarian TIME are associated with worse prognosis (10,46,47), our findings show a spatial and context dependency on CD163-mediated activities. Therapeutically, CD163 targeting strategies (e.g., OR2805) have shown to be effective in relieving immune suppression and are therefore clinically evaluated in a trial for solid tumors (48), thus representing an opportunity to target the robust HGSC TIME-associated immune suppressive macrophages to potentially improve anti-tumor immune surveillance (49,50).

While our results partially align with previous studies, for example in the finding for B cell-M1 macrophage interactions, we did not achieve significance in univariate correlations to patient outcomes for T cell and macrophage proportions as expected. While we did find Hazard Ratio estimates in the expected direction (Hazard Ratio estimates for infiltration by all T cell populations and M1 macrophages were <1), our results were not significant. Notably, increased CD8+ T cells conveyed a Hazard Ratio of 0.74 (p=0.059) and CD8+ T cell proportion (HR 0.68, p=0.035) and CD4+ T cell proportion (HR 0.58, p=0.02) were significantly correlated with improved OS for only the primary tumor samples. The vascular endothelial cell proportion, M1 macrophage proportion, and CD4+ T cell proportion were also chosen by the random forest as important features for predicting PFS. We emphasize the importance of differences in the definitions of cell types when comparing our results with previous works. For example, prior literature has suggested that M2 macrophages are typically more prevalent than M1 macrophages in HGSC (51), which contrasts with our results (Fig. 2A). However, if we had included CD163+ cells in the M2 macrophage cluster (52,53), then the M2 macrophage count would indeed be higher than the M1 macrophage count alone and present findings in line with the aforementioned study. An explanation for differences with previous studies might be due to differences in cell clustering and phenotyping, pointing to the need for further refinement of consistent markers, particularly so that such results can become relevant in clinical application.

Limitations of the imaging technology used in this study affect the significance of our findings. In particular, the FOV size of 500 μm at single-cell resolution might still be a limiting factor for the comprehensive documentation of the clinically relevant spatial organization in the TIME. Despite staining with antibodies to 26 proteins, an average of 14% of cells remained unidentified in the 77 samples included in our final analysis, due to them not expressing any of the phenotypic markers. The spatial organization of the TIME may be better delineated in a more comprehensive higher parameter analysis tailored to identification of cells in HGSC. For example, future work might additionally use functional markers to further characterize CD163+ cells. In our analyses we treated the set of unidentified cells as a population and found that they contributed to significant interactions, highlighting an opportunity for future research. For example, the contact enrichment score between unidentified cells and M1 macrophages was significantly associated with worse PFS (HR=1.41, p=0.01) and chosen as one of the most important features for predicting both OS and PFS.

Our study investigated the relative importance of different types and combinations of clinical/immunohistochemical and TIME features in modeling patient outcomes before treatment via both feature importance values within a random forest model for out-of-sample prediction and coefficient values within Cox regressions on in-sample data. While one tumor sample was from 1996, and aspects of clinical management have improved over the time period during which the samples were generated (e.g., increased testing for BRCA mutation), we assume that better prognosis in this dataset largely is due to differential response to a standard of care treatment, which has not changed substantially since 1996. Cox regressions evaluated on in-sample data can be used to describe observed patterns, but do not provide results about out-of-sample predictive performance relevant for generalizing our results to new patients in clinical contexts. We also primarily report results from univariate analyses which only consider features in isolation and multivariate Cox regressions with all significant features did not converge. Due to the exploratory nature of the study, we report non-adjusted p-values, and we found no significant univariate correlations with false discovery rate adjusted p-values (33).

Although random forests are a popular choice in predictive modeling, in part because of their built-in regularization controls for overfitting and their strong interpretability (34), all machine learning models are potentially vulnerable to overfitting. In our analysis, we did not observe substantially better-than-random out of sample predictive performance on patient OS on average, indicating that the features chosen as relatively important for predicting OS might have been used by the model to overfit (i.e., learn complex rules to fit to the training dataset that do not generalize to predictive performance on unseen data), and thus might be considered with more skepticism than those chosen as relatively important for predicting PFS. We took care to avoid cases in which the data used to evaluate or test the model’s accuracy was not fully independent of the data used to train the model, for example by imputing NA values separately in the train and test sets, which is one cause of overfitting. At the same time, our training data were derived from 77 of patients whose corresponding feature sets may not be fully representative of the underlying biology, implying that the reported predictive accuracies should be interpreted cautiously, and more weight should be placed on the inference that some features are relatively more important than others in the prediction of patient outcomes.

In comparing categories of features based on their respective processes of derivation, we found that models including features derived from spatial network representations of the TIME performed slightly worse. However, our results do support the continued use of spatial networks in quantifying and evaluating the TIME. Network features were the largest and most diverse category of features we evaluated (N=117), and many of them were irrelevant for predicting patient outcomes, as indicated by low Gini importance scores, thus likely reducing predictive performance for the network feature category as a whole by introducing noise. However, feature importance evaluations indicate that a subset of these network features were among some of the most important features overall for prediction: in particular, the contact enrichment features encoding information about the proximity between cell types were generally ranked as more important than mean region size or assortativity features, which encode cell clustering patterns (Fig. S11, Fig. S12), mirroring a similar finding in the in-sample Cox regression results. Further development and refinement of features derived from spatial network representations of the TIME could potentially improve the development of useful markers.

Many of the features identified as important for patient outcome prediction involved the spatial relationship between cells other than tumor cells. While not surprising, this finding strongly emphasizes the importance of investigating cell-cell interactions throughout the TIME. Two overarching goals of such studies would be to (i) identify key cell types that can be directly addressed with targeted therapies, and (ii) to develop methods that more generally help to characterize the TIME prior to patient treatment. For instance, further studies might investigate how the spatial organization of the TIME differs between tumor sites within the same patient, and whether this can drive differential response to treatment between tumor sites (54). Additionally, future work could build on previous studies investigating TIME changes with chemotherapy treatment (55,56) to investigate how the spatial organization of the TIME changes with chemotherapy. Those goals aim to improve individualized patient diagnosis and care while at the same time enhancing our understanding of more general pathways of cancer development and progression.

## MATERIALS AND METHODS

### Study design

We procured formalin-fixed paraffin-embedded tumor samples from patients diagnosed with HGSC of the ovary, fallopian tube, and peritoneum under the University of Colorado’s IRB Protocol, COMIRB #17-7788. The tumor samples were examined by a Gynecologic Pathologist (Dr. Miriam Post) and viable tumor areas were selected for generation of the tissue microarray. The total number of tumors on the tissue microarray was 133, which include primary and recurrent HGSC tumors. Further details of the tissue microarray can be found in Watson et al. 2019 (18), Jordan et al. 2020 (19), and McMellen et al. 2023 (20). Multiplexed ion beam imaging was performed on 83 tumor specimens. All samples were from patients with cancer of ovary, fallopian tube, and peritoneum diagnosed at a similar stage (IIIC). For 77 (69 primary and 8 recurrent) of these samples with sufficient cell type identification to produce spatial features (see Fig. S1), we studied clinical/immunohistochemical features in combination with descriptive (composition, spatial, and network features) features derived from these samples. The study design aims to integrate features that could hypothetically be generated from a patient’s biopsy samples before treatment in an exploratory analysis to investigate what features or combination of features could be used to predict patient outcomes and motivate adjustments in treatment.

### Clinical/immunohistochemical features

For each sample, we investigated six clinical/immunohistochemical features alone and in combination with features derived from the samples: BRCA mutational status, age, and histology scores for H3K14Ace status, ATF6 status, DUSP1 status, and CBX2 status, calculated by multiplying the intensity of the stain [0-3] by the percentage of that intensity [0-100].

BRCA mutational status was included because of its well-established risk and therapeutic implication (21,22). The remaining features were selected and included based on prior work (20,23,24) that demonstrated prognostic value. Age was included because it is a prognostication indicator in terms of OS (25). Figure S7 shows distributions of all the clinical/immunohistochemical features across the final 77 samples analyzed.

### MIBI-TOF imaging

Imaging was performed using a custom MIBI-TOF instrument with a Xe^+^ primary ion source upgrade (17). A total of 83 images with a field of view size of 500×500 µm and a frame size of 1024×1024 pixels were acquired. The beam current was set to 5 nA with a dwell time of 2 ms, yielding a resolution of approximately 0.5 µm per pixel. Secondary ions were accelerated into the time-of-flight mass spectrometer with a sample bias of 50 V and detected with a temporal resolution of 0.6 ns across a mass range of 1-200 m/z^+^.

### Low-level image processing

Multiplexed images were extracted and processed using Ionpath’s MIBI/O software: The image data was background- and mass-corrected with vendor-provided configuration files, see Table S1. In the next step, the image data was denoised with the filtering parameters provided in File D2.

### Low-level image pre-processing

We adopted a custom computational pipeline developed to analyze MIBI data (26). In this framework multi-step low-level image processing is replaced with a single-step pixel classification where each pixel in an image is classified such that all categories of undesired signal are placed in a different class from the desired marker signal and continue the downstream analysis using the generated feature representation map of the marker signal.

### Single-cell segmentation

Whole-cell segmentation was done using the pre-trained single-cell segmentation model Mesmer (27). We used the dsDNA channel for nuclear segmentation and the β-tubulin channel to guide identification of cell boundaries.

### Cell-type identification

Single-cell data were extracted for all the cells and normalized by the cell size. To assign each cell to a lineage, we used the unsupervised clustering algorithm as implemented in FlowSOM (28) with multiple steps: first we identified the immune cells and non-immune cells using the following markers: CD45, HLA-DR, CD31, Podoplanin, Vimentin, and Keratin. Then, we used the immune markers CD3, CD4, CD8, CD20, CD68, CD56, CD11b, CD11c, CD163, DC-SIGN to identify the immune subsets (See Table S6).

### Spatial and network features

We calculated spatial and network features from the sample images following Moldoveanu et al. 2022 (16) using Python version 3.9.12, *SciPy* version 1.7.3 and *NetworkX* version 2.7.1. We calculated the median Euclidean distance in pixels in each sample between cells of three focal cell types: tumor cells, M1 macrophages, and vascular endothelial cells and their nearest neighbor of all other non-focal cell types. In each sample, we examined each of the focal cells of interest and then identified the nearest neighbor of each non-focal cell type using KD Trees (implemented in *SciPy*) and recorded the Euclidean distances. For each sample, we report the median nearest neighbor distance for each combination of non-focal cell type (listed first) and focal cell type.

Spatial network representations of the samples were created by connecting spatial neighbors identified using Delaunay triangulation (implemented in *SciPy*) and then trimming edges that were above a threshold of 50 pixels (∼24.4 μm). Results were not sensitive to using a higher threshold for trimming edges (100 pixels, ∼48.8 μm, Fig. S13-15). Versions of the spatial networks were created in which neighboring cells were only connected if they were of the same cell type and connected regions of the same cell type were identified from these modified networks using the `connected_components` function implemented in *NetworkX*. The mean of the region sizes in each sample were reported for each cell type.

Binary attributes were added to each cell in the spatial networks for each cell type, set to 1 if the cell was of that type and 0 otherwise. Assortativity coefficients (29) were then calculated using the *NetworkX* function `attribute_assortativity_coefficient` for each of these binary cell type attributes, thus measuring to what extent cells tended to be neighbors with cells of the same type versus any other type. This value is 1 for perfect assortative mixing, in which cells are only neighbors with cells of the same type, 0 when there is no assortative mixing, and negative when there is disassortative mixing, in which cells are typically neighbors with cells of different types.

We calculated contact enrichment scores for the three focal cell types of tumor cells, M1 macrophages, and vascular endothelial cells and each non-focal cell type. Following a procedure used in prior work (16,30,31), the cell type labels of all cells other than those of the focal cell types were randomized 1000 times. After each shuffle, the number of times that the focal cell type was a neighbor of each non-focal cell type in the spatial network is recorded. These counts represent a null distribution for each non-focal cell type which is then compared to the observed number of contacts, and the z-score is recorded as the contact enrichment score. A negative contact enrichment score thus indicates fewer contacts than expected at random, a contact enrichment score of 0 indicates as many, and a positive contact enrichment score indicates more contacts than expected at random. When a cell type was missing from a sample, we recorded the mean region, contact enrichment and assortativity values as 0 and the median nearest neighbor distances as “NA” for features related to that cell type for the sample. We also report results for both the Cox regression and random forest analyses in the supplementary material when recording these values all as “NA” (See Note S1, File D8, Fig. S16-18).

### Statistical analysis

We fit Cox proportional hazards regression models (32) to OS and PFS outcomes using the `coxph` function from the *survival* package (version 3.5-5) in R (version 4.3.1). Univariate regressions were performed with each of the 216 clinical/immunohistochemical, composition, spatial, and network features treated as individual covariates. All covariates except BRCA Mutation and age were z-score normalized before analysis so that coefficients were comparable across different feature scales and any rows with NA values were excluded. A covariate was considered significant if it had a p-value of under 0.05. In the Supplementary Materials (File D5) we report the number of samples considered for each regression, the number of relevant events considered in the time to event analysis (death or recurrence, respectively), the covariate’s coefficient in the Cox proportional hazard regression, the corresponding hazard ratio, and the p-value and false discovery rate adjusted p-value (33). Given the exploratory nature of this study, we focused on results that were significant with non-adjusted p-values. Multivariate Cox proportional hazards models were fitted using the `coxphmulti` function. Models with all covariates found to be significant in univariate regressions for both outcome variables did not converge, so we ran multivariate models with the top five features for each outcome variable, ranked by p-value, both adjusted for clinical/immunohistochemical attributes and as a reduced model without an adjustment for clinical/immunohistochemical attributes (Table S2, S3).

### Predictive analysis

We used a random forest classification model (34) implemented in the R package *randomForest* (version 4.7-1.1), R version 4.3.1 with default hyperparameters (see Note S2). Random forests were chosen as our predictive method because they have been shown to work well on high-dimensional data with a low sample size and can be used to rank features based on importance scores (35).

We first investigated what subsets of features, based on all possible combinations of the four feature categories (clinical/immunohistochemical, composition, spatial, and network features), produced the highest expected out-of-sample predictive performance: We repeated 500 classification tasks for each of the two outcome variables and 15 models. For each of these classification tasks, 70% of the samples were treated as a training set and 30% were treated as a test set. Data was split randomly for each classification task using the `sample.split` function in the package *caTools* (version 1.18.2) in order to preserve the ratio between outcome labels in the two sets. NA values were imputed separately in the training and test set using the `na.roughfix` function from the *randomForest* package which performs median substitution for numeric variables and mode substitution for factor variables. In the rare cases when a train and test split were selected such that a feature was entirely NA in the test set, we did not use that train-test split and instead re-drew. Predictive performance was calculated for each classification task using the AUC (Area under the receiver operating characteristics curve) statistic, calculated using the `roc` and `auc` functions in the *pROC* package, with the direction parameter set such that positive samples should receive a higher predicted value (version 1.18.2). The AUC statistic was chosen because of its properties of being threshold invariant, scale invariant, and use-case agnostic, hence providing a useful measure by which to compare the general performance of different models (36).

Second, we investigated an overall ranking of feature importance from the models including all features. Features were ranked based on their median Gini importance across the 500 classification tasks. The Gini importance for a feature indicates the mean decrease in node impurity caused by splitting on that feature during model training, in which higher values indicate that the feature was more useful during the generation of the random forest model. The Gini importance can be biased to provide higher importance values for numeric features as they exhibit more potential split points (37). However, this bias would not have a strong influence on our results because the BRCA mutation status is the only categorical variable in our dataset. Repeating the evaluation 500 times allowed us to explore consistency and variation in the ranking of the features across different train and test splits of the data, which we chose to do based on the small sample size and the expectation that many of the generated features would be highly correlated.

## Acknowledgements

We acknowledge philanthropic contributions from Kay L. Dunton Endowed Memorial Professorship In Ovarian Cancer Research, the McClintock-Addlesperger Family, Karen M. Jennison, Don and Arlene Mohler Johnson Family, Michael Intagliata, Duane and Denise Suess, Mary Normandin, and Donald Engelstad. We thank Samreen Anjum for conversations about the predictive modeling. We thank Dr. Angelo’s laboratory at Stanford for assistance with MIBI instrument setup and recording time. This work was supported by The American Cancer Society (B Bitler, RSG-19-129-01-DDC), NIH/NCI (B Bitler, R37CA261987 and R01CA285446), and Ovarian Cancer Research Alliance (A Clauset and B Bitler), and the University of Colorado Cancer Center Support Grant (P30CA046934). We also thank the Human Immune Monitoring Shared Resource (RRID:SCR_021985) for their expert assistance with sample preparation and data acquisition.

## Informed consent statement

A TMA comprised of serous tumors (COMIRB #17-7788) was used. This protocol is deemed exempt, as it is using previously collected data, and the information is not recorded in a manner that is identifiable. Further, the findings of the study did not alter treatment choices or patient outcomes.

## Data availability statement

Data and code used to perform analyses and supplemental data files are available upon request.

## References

1. Lisio M-A, Fu L, Goyeneche A, Gao Z, Telleria C. High-Grade Serous Ovarian Cancer: Basic Sciences, Clinical and Therapeutic Standpoints. IJMS. 2019;20:952.

2. Siegel RL, Miller KD, Wagle NS, Jemal A. Cancer statistics, 2023. CA A Cancer J Clinicians. 2023;73:17–48.

3. Hoppenot C, Eckert MA, Tienda SM, Lengyel E. Who are the long-term survivors of high grade serous ovarian cancer? Gynecologic Oncology. 2018;148:204–12.

4. Garsed DW, Pandey A, Fereday S, Kennedy CJ, Takahashi K, Alsop K, et al. The genomic and immune landscape of long-term survivors of high-grade serous ovarian cancer. Nat Genet. 2022;54:1853–64.

5. Coscia F, Lengyel E, Duraiswamy J, Ashcroft B, Bassani-Sternberg M, Wierer M, et al. Multi-level Proteomics Identifies CT45 as a Chemosensitivity Mediator and Immunotherapy Target in Ovarian Cancer. Cell. 2018;175:159–170.e16.

6. Launonen I-M, Lyytikäinen N, Casado J, Anttila EA, Szabó A, Haltia U-M, et al. Single-cell tumor-immune microenvironment of BRCA1/2 mutated high-grade serous ovarian cancer. Nat Commun. 2022;13:835.

7. Kandalaft LE, Odunsi K, Coukos G. Immune Therapy Opportunities in Ovarian Cancer. American Society of Clinical Oncology Educational Book. 2020;e228–40.

8. Zhang L, Conejo-Garcia JR, Katsaros D, Gimotty PA, Massobrio M, Regnani G, et al. Intratumoral T Cells, Recurrence, and Survival in Epithelial Ovarian Cancer. N Engl J Med. 2003;348:203–13.

9. Nielsen JS, Sahota RA, Milne K, Kost SE, Nesslinger NJ, Watson PH, et al. CD20+ Tumor-Infiltrating Lymphocytes Have an Atypical CD27− Memory Phenotype and Together with CD8+ T Cells Promote Favorable Prognosis in Ovarian Cancer. Clinical Cancer Research. 2012;18:3281–92.

10. Yuan X, Zhang J, Li D, Mao Y, Mo F, Du W, et al. Prognostic significance of tumor-associated macrophages in ovarian cancer: A meta-analysis. Gynecologic Oncology. 2017;147:181–7.

11. Chen J, Wang Y, Ko J. Single-cell and spatially resolved omics: Advances and limitations. Journal of Pharmaceutical Analysis. 2023;13:833–5.

12. Williams CG, Lee HJ, Asatsuma T, Vento-Tormo R, Haque A. An introduction to spatial transcriptomics for biomedical research. Genome Med. 2022;14:68.

13. Fang S, Chen B, Zhang Y, Sun H, Liu L, Liu S, et al. Computational Approaches and Challenges in Spatial Transcriptomics. Genomics, Proteomics & Bioinformatics. 2023;21:24–47.

14. Lee J, Yoo M, Choi J. Recent advances in spatially resolved transcriptomics: challenges and opportunities. BMB Rep. 2022;55:113–24.

15. Steinhart B, Jordan KR, Bapat J, Post MD, Brubaker LW, Bitler BG, et al. The Spatial Context of Tumor-Infiltrating Immune Cells Associates with Improved Ovarian Cancer Survival. Molecular Cancer Research. 2021;19:1973–9.

16. Moldoveanu D, Ramsay L, Lajoie M, Anderson-Trocme L, Lingrand M, Berry D, et al. Spatially mapping the immune landscape of melanoma using imaging mass cytometry. Sci Immunol. 2022;7:eabi5072.

17. Greenbaum S, Averbukh I, Soon E, Rizzuto G, Baranski A, Greenwald NF, et al. A spatially resolved timeline of the human maternal–fetal interface. Nature. 2023;619:595– 605.

18. Watson ZL, Yamamoto TM, McMellen A, Kim H, Hughes CJ, Wheeler LJ, et al. Histone methyltransferases EHMT1 and EHMT2 (GLP/G9A) maintain PARP inhibitor resistance in high-grade serous ovarian carcinoma. Clin Epigenet. 2019;11:165.

19. Jordan KR, Sikora MJ, Slansky JE, Minic A, Richer JK, Moroney MR, et al. The Capacity of the Ovarian Cancer Tumor Microenvironment to Integrate Inflammation Signaling Conveys a Shorter Disease-free Interval. Clinical Cancer Research. 2020;26:6362–73.

20. McMellen A, Yamamoto TM, Qamar L, Sanders BE, Nguyen LL, Ortiz Chavez D, et al. ATF6-Mediated Signaling Contributes to PARP Inhibitor Resistance in Ovarian Cancer. Molecular Cancer Research. 2023;21:3–13.

21. Gori S, Barberis M, Bella MA, Buttitta F, Capoluongo E, Carrera P, et al. Recommendations for the implementation of BRCA testing in ovarian cancer patients and their relatives. Critical Reviews in Oncology/Hematology. 2019;140:67–72.

22. Moschetta M, George A, Kaye SB, Banerjee S. BRCA somatic mutations and epigenetic BRCA modifications in serous ovarian cancer. Annals of Oncology. 2016;27:1449–55.

23. Sanders BE, Yamamoto TM, McMellen A, Woodruff ER, Berning A, Post MD, et al. Targeting DUSP Activity as a Treatment for High-Grade Serous Ovarian Carcinoma. Molecular Cancer Therapeutics. 2022;21:1285–95.

24. Wheeler LJ, Watson ZL, Qamar L, Yamamoto TM, Post MD, Berning AA, et al. CBX2 identified as driver of anoikis escape and dissemination in high grade serous ovarian cancer. Oncogenesis. 2018;7:92.

25. Peres LC, Sinha S, Townsend MK, Fridley BL, Karlan BY, Lutgendorf SK, et al. Predictors of survival trajectories among women with epithelial ovarian cancer. Gynecologic Oncology. 2020;156:459–66.

26. Ahmadian M, Rickert C, Minic A, Wrobel J, Bitler BG, Xing F, et al. A platform-independent framework for phenotyping of multiplex tissue imaging data. Uhlmann V, editor. PLoS Comput Biol. 2023;19:e1011432.

27. Greenwald NF, Miller G, Moen E, Kong A, Kagel A, Dougherty T, et al. Whole-cell segmentation of tissue images with human-level performance using large-scale data annotation and deep learning. Nat Biotechnol. 2022;40:555–65.

28. Van Gassen S, Callebaut B, Van Helden MJ, Lambrecht BN, Demeester P, Dhaene T, et al. FlowSOM: Using self-organizing maps for visualization and interpretation of cytometry data. Cytometry Pt A. 2015;87:636–45.

29. Newman MEJ. Mixing patterns in networks. Phys Rev E. 2003;67:026126.

30. Keren L, Bosse M, Marquez D, Angoshtari R, Jain S, Varma S, et al. A Structured Tumor-Immune Microenvironment in Triple Negative Breast Cancer Revealed by Multiplexed Ion Beam Imaging. Cell. 2018;174:1373–1387.e19.

31. Schapiro D, Jackson HW, Raghuraman S, Fischer JR, Zanotelli VRT, Schulz D, et al. histoCAT: analysis of cell phenotypes and interactions in multiplex image cytometry data. Nat Methods. 2017;14:873–6.

32. Andersen PK, Gill RD. Cox’s Regression Model for Counting Processes: A Large Sample Study. Ann Statist [Internet]. 1982 [cited 2023 Jul 31];10. Available from: https://projecteuclid.org/journals/annals-of-statistics/volume-10/issue-4/Coxs-Regression-Model-for-Counting-Processes--A-Large-Sample/10.1214/aos/1176345976.full

33. Benjamini Y, Hochberg Y. Controlling the False Discovery Rate: A Practical and Powerful Approach to Multiple Testing. Journal of the Royal Statistical Society: Series B (Methodological). 1995;57:289–300.

34. Breiman L. Random Forests. Machine Learning. 2001;45:5–32.

35. Chen X, Ishwaran H. Random forests for genomic data analysis. Genomics. 2012;99:323–9.

36. Fawcett T. An introduction to ROC analysis. Pattern Recognition Letters. 2006;27:861–74.

37. Nembrini S, König IR, Wright MN. The revival of the Gini importance? Valencia A, editor. Bioinformatics. 2018;34:3711–8.

38. Macciò A, Gramignano G, Cherchi MC, Tanca L, Melis L, Madeddu C. Role of M1-polarized tumor-associated macrophages in the prognosis of advanced ovarian cancer patients. Sci Rep. 2020;10:6096.

39. Rubatt JM, Darcy KM, Hutson A, Bean SM, Havrilesky LJ, Grace LA, et al. Independent prognostic relevance of microvessel density in advanced epithelial ovarian cancer and associations between CD31, CD105, p53 status, and angiogenic marker expression: A Gynecologic Oncology Group study. Gynecologic Oncology. 2009;112:469–74.

40. Pinto MP, Balmaceda C, Bravo ML, Kato S, Villarroel A, Owen GI, et al. Patient inflammatory status and CD4+/CD8+ intraepithelial tumor lymphocyte infiltration are predictors of outcomes in high-grade serous ovarian cancer. Gynecologic Oncology. 2018;151:10–7.

41. Mhawech-Fauceglia P, Wang D, Ali L, Lele S, Huba MA, Liu S, et al. Intraepithelial T cells and tumor-associated macrophages in ovarian cancer patients. Cancer Immunity. 2013;13.

42. Lieber S, Reinartz S, Raifer H, Finkernagel F, Dreyer T, Bronger H, et al. Prognosis of ovarian cancer is associated with effector memory CD8 ^+^ T cell accumulation in ascites, CXCL9 levels and activation-triggered signal transduction in T cells. OncoImmunology. 2018;7:e1424672.

43. Mantovani A, Marchesi F, Malesci A, Laghi L, Allavena P. Tumour-associated macrophages as treatment targets in oncology. Nat Rev Clin Oncol. 2017;14:399–416.

44. Zhang M, He Y, Sun X, Li Q, Wang W, Zhao A, et al. A high M1/M2 ratio of tumor-associated macrophages is associated with extended survival in ovarian cancer patients. 2014;

45. Buechler C, Ritter M, Orsó E, Langmann T, Klucken J, Schmitz G. Regulation of scavenger receptor CD163 expression in human monocytes and macrophages by pro- and antiinflammatory stimuli. Journal of Leukocyte Biology. 2000;67:97–103.

46. No JH, Moon JM, Kim K, Kim Y-B. Prognostic Significance of Serum Soluble CD163 Level in Patients with Epithelial Ovarian Cancer. Gynecol Obstet Invest. 2013;75:263–7.

47. Lan C, Huang X, Lin S, Huang H, Cai Q, Wan T, et al. Expression of M2-Polarized Macrophages is Associated with Poor Prognosis for Advanced Epithelial Ovarian Cancer. Technol Cancer Res Treat. 2013;12:259–67.

48. OncoResponse, Inc. A Phase 1-2 Study of OR2805, a Monoclonal Antibody Targeting CD163, Alone and in Combination With Anticancer Agents in Subjects With Advanced Malignancies [Internet]. clinicaltrials.gov; 2022 Sep. Report No.: NCT05094804. Available from: https://clinicaltrials.gov/study/NCT05094804

49. Chen T, Chen J, Zhu Y, Li Y, Wang Y, Chen H, et al. CD163, a novel therapeutic target, regulates the proliferation and stemness of glioma cells via casein kinase 2. Oncogene. 2019;38:1183–99.

50. Probst P, Simmons R, Dinh H, Zuck M, Wall V, Bouchlaka M, et al. 271 Development of OR2805, an anti-CD163 antibody derived from an elite responder to checkpoint inhibitor therapy that relieves immunosuppression caused by M2c macrophages. J Immunother Cancer. 2021;9:A294–A294.

51. Schweer D, McAtee A, Neupane K, Richards C, Ueland F, Kolesar J. Tumor-Associated Macrophages and Ovarian Cancer: Implications for Therapy. Cancers. 2022;14:2220.

52. Biswas SK, Mantovani A. Macrophage plasticity and interaction with lymphocyte subsets: cancer as a paradigm. Nat Immunol. 2010;11:889–96.

53. Sulahian TH, Högger P, Wahner AE, Wardwell K, Goulding NJ, Sorg C, et al. Human monocytes express CD163, which is upregulated by IL-10 and identical to p155. Cytokine. 2000;12:1312–21.

54. Masoodi T, Siraj S, Siraj AK, Azam S, Qadri Z, Parvathareddy SK, et al. Genetic heterogeneity and evolutionary history of high-grade ovarian carcinoma and matched distant metastases. Br J Cancer. 2020;122:1219–30.

55. Mesnage SJL, Auguste A, Genestie C, Dunant A, Pain E, Drusch F, et al. Neoadjuvant chemotherapy (NACT) increases immune infiltration and programmed death-ligand 1 (PD-L1) expression in epithelial ovarian cancer (EOC). Annals of Oncology. 2017;28:651–7.

56. Cao G, Hua D, Li J, Zhang X, Zhang Z, Zhang B, et al. Tumor immune microenvironment changes are associated with response to neoadjuvant chemotherapy and long-term survival benefits in advanced epithelial ovarian cancer: A pilot study. Front Immunol. 2023;14:1022942.

