## Supplementary Material for "The spatial structure of the tumor immune microenvironment can explain and predict patient response in high-grade serous carcinoma"

**Table of Contents**

**Supplementary Materials and Methods**

Note S1. Handling missing cell types in the analysis

Note S2. Random forest hyperparameter selection sensitivity tests

**Supplementary Figures**

Figure S1. Exclusion criteria for samples excluded from the analysis

Figure S2. Spatial network creation demonstration

Figure S3. Median neighbor distances in the spatial networks

Figure S4. Cell type region size distributions

Figure S5. Survival curves

Figure S6. Outcome variable distributions

Figure S7. Clinical/immunohistochemical feature distributions

Figure S8. Univariate Cox regression results for only primary tumor samples

Figure S9. Radom forest predictive performance results for only primary tumor samples

Figure S10. Random forest feature importance results for only primary tumor samples

Figure S11. Random forest feature importance results split by feature type - OS

Figure S12. Random forest feature importance results split by feature type - PFS

Figure S13. Univariate Cox regression results with increased spatial network trimming threshold

Figure S14. Random forest predictive performance results with increased spatial network trimming threshold

Figure S15. Random forest feature importance results with increased spatial network trimming threshold

Figure S16. Univariate Cox regression results with features derived from missing cell types all treated as NA

Figure S17. Random forest predictive performance results with features derived from missing cell types all treated as NA

Figure S18. Random forest feature importance results with features derived from missing cell types all treated as NA

### Supplementary Tables

Table S1. Mass correction parameters

Table S2. Multivariate Cox regression results - OS

Table S3. Multivariate Cox regression results - PFS

Table S4. Random forest predictive performance results

Table S5. Random forest predictive performance results for only primary tumor samples

Table S6. Definitions of cellular phenotypes identified with unsupervised clustering

### Supplementary Data Files

The following supplementary data files are available upon request to the corresponding authors:

D1. Antibody panel design

D2. Noise filtering parameters for image data

D3. Correlation coefficients between other cell type percentages and tumor cell percentages

D4. Univariate Cox regression results for only primary tumor samples

D5. Univariate Cox regression main text results

D6. Ranking of feature importance in the random forest model for only primary tumor samples

D7. Ranking of feature importance in the random forest model for main text results

D8. Univariate Cox regression results with features derived from missing cell types all treated as NA

D9. Results of 5-fold cross-validation results with hyperparameter selection

**Supplementary Materials and Methods:**

**Note S1. Handling missing cell types in the analysis**

Our analysis involved generating a number of features based on the composition and spatial structure of the samples. Not every identified cell type was present in all samples. For example, dendritic cells were relatively rare in our dataset, only identified in 23 of the 77 samples. We explored handling these missing cell types in two different ways when generating features for a sample.

In the first way, corresponding to the results presented in the main text (Fig.5-7, File D4, D6, D7, Table S2, S3), we filled in these missing values based on knowledge of the relevant feature type. Cell type proportion values and mean region sizes were recorded as 0 if the cell type was missing from the sample given that these are size values. Contact enrichment scores between a missing cell type and tumor cells, M1 macrophages, or vascular endothelial cells were also recorded as 0. Because contact enrichment scores are z-scores, a value of 0 indicates that the number of contacts between this missing cell type and the focal cell type does not differ from null expectations, which would be 0 contacts. Assortativity coefficient values were also recorded as 0 for missing cell types, or in cases in which there was only one cell of a given type in the sample, given that the assortativity coefficient takes on the value of 0 when there is neither assortative or disassortative mixing indicated (1). The median nearest neighbor distances to tumor cells, M1 macrophages, and vascular endothelial cells were recorded as NA for missing cell types, as setting this value to 0 would imply that the missing cell type was unexpectedly close to the focal cell type in the sample. We also recorded as 0 all of the contact enrichment scores and as NA all of the median nearest neighbor distance values with vascular endothelial cells or M1 macrophages as the focal cell type for samples in which vascular endothelial cells or M1 macrophages were missing from the sample.

In the second approach, corresponding to the results presented in Fig. S10-S12 (File D8, D9), we left all derived features related to missing cell types as NA. We present results using the first approach in the main text because it uses knowledge of the feature types.

We observed similar predictive performances across models using the two approaches, with slightly higher AUC scores, still under the 0.5 threshold, for OS using the approach displayed in Figure S17. We present these results because, while many results were not sensitive to this data processing choice, the composition and ranking of significant features in the Cox regressions and the top ten features in the predictive models do slightly change, so those features that are not consistently identified between the two data processing choices should be regarded with more caution when developing future hypotheses.

With both data processing choices, remaining NA values were handled in the same way before running the Cox regression and random forest analyses. In the Univariate Cox regressions, we excluded samples with NA values for a given covariate, thus generally running regressions on fewer samples to produce the results shown in Figure S16. In the random forest analysis, NA values were imputed separately in the training and test set using median substitution for numeric variables and mode substitution for factor variables before model training and evaluation.

**Note S2. Random forest hyperparameter selection sensitivity tests**

The reported results in the main text use the default values in the R randomForest package for the *nodesize* and *mtry* hyperparameters. The *nodesize* hyperparameter has a default value of 1 for classification and represents the minimum size of terminal nodes in the tree, and thus can be used to build smaller trees if the value is increased. The *mtry* hyperparameter has a default value of $\sqrt{p}$ (where $p$ is the number of features) for classification and represents the number of features selected as candidates at each split, which can produce more diverse trees with a higher value. We tested the sensitivity of predictive performance results on this dataset to hyperparameter selection by reporting average AUC results across folds (2) from 5-fold cross validation with hyperparameter selection for each of the 15 models (File D7). Hyperparameters were selected via 5-fold cross-validation on each training set and a grid search (*nodesize* ranging 1-20 and *mtry* ranging 2-12), where the best combination of the two parameters for the training set was selected based on the AUC and then used to evaluate predictive performance on the held out fold.

We found that the average predictive performance results of the 15 models after hyperparameter selection were similar to those obtained averaging performance across 500 evaluations using default hyperparameters, only slightly improving performance in some cases. The best model for PFS, model 8, for example, achieved an average AUC of 0.745 across cross-validation folds with hyperparameter selection, improving from an average AUC of 0.711 using default hyperparameters, and models 8, 11, and 3 were still the top models for predicting PFS. The optimal hyperparameters chosen tended to opt for smaller trees (larger nodesize values), which likely slightly improved model performance by decreasing overfitting. We did not perform hyperparameter selection for each of the 500 evaluations used to produce results in the main text due to the computational intensity of running a 10x20 hyperparameter grid search for 500 evaluations each of 30 models (as opposed to 5 evaluations each of 30 models in the 5-fold cross validation test).

**Supplementary Figures:**

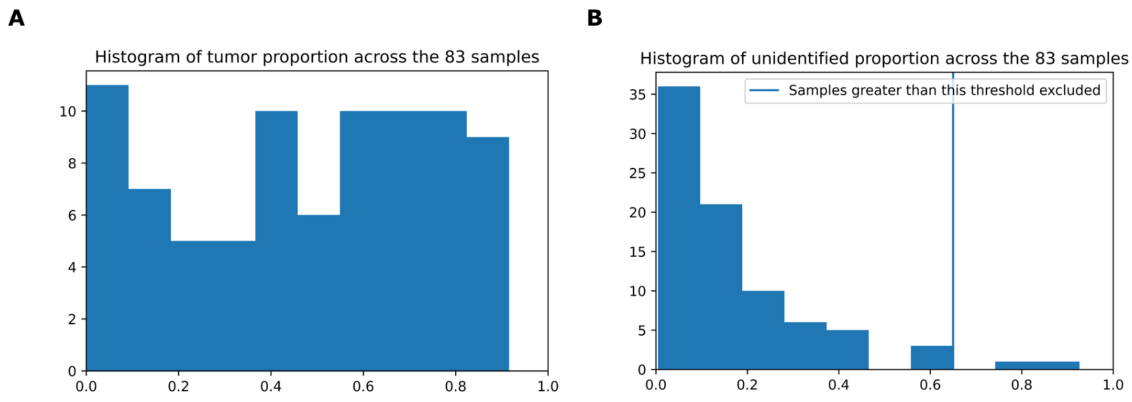


**Fig S1. Exclusion criteria for samples excluded from the analysis.** Samples were excluded from the analysis based on cell type proportions. (A) tumor cell proportion across the samples, samples were only excluded if they had no tumor cells (N=3). (B) Unidentified cell proportion across the samples, samples were excluded if their unidentified cell proportion was over 0.65 (N=2).

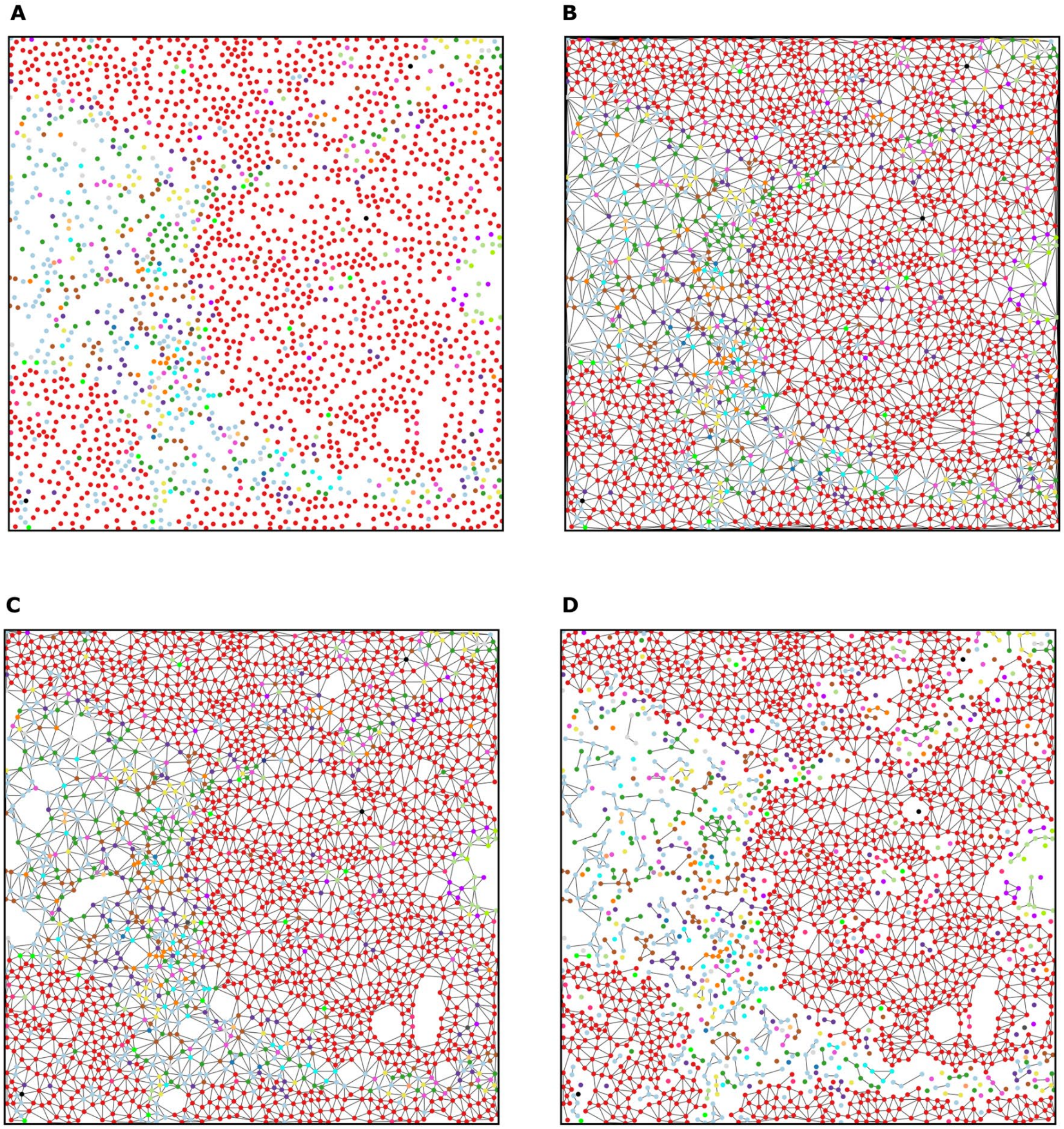

**Fig S2. Spatial network creation demonstration.** (A) One of the samples (ID 9) visualized after cell segmentation and phenotyping, with each cell type a different color. (B) First, Delaunay triangulation was used to connect spatial neighbors. (C) Next, the edges larger than 50 pixels (~24.4 μm) were removed to create the final spatial network used in the analysis. (D) To compute the mean region size, edges between cells of different types were removed and we computed the mean size of connected components for each cell type.

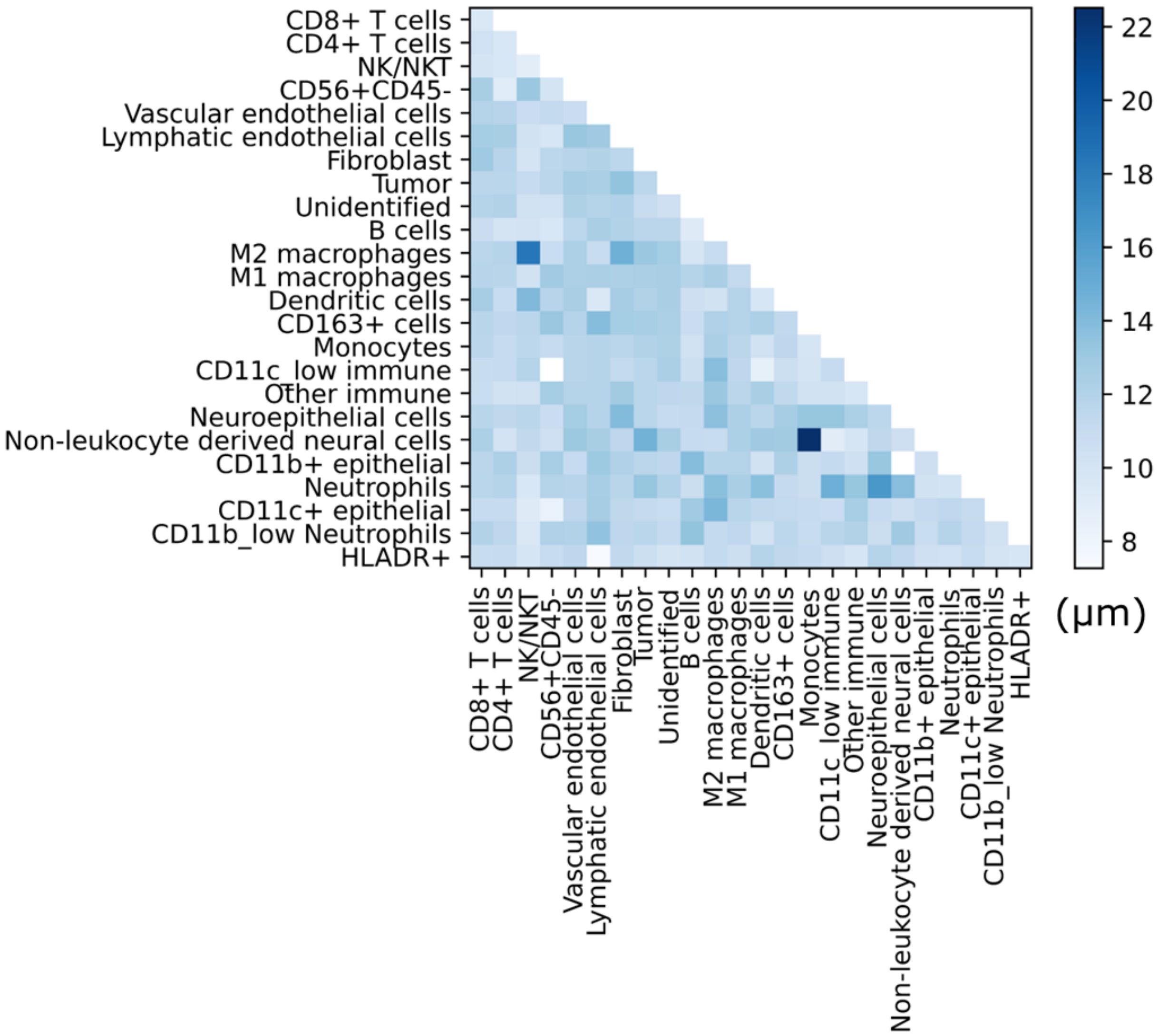


**Fig S3. Median neighbor distances in the spatial networks.** Heatmap of the median distances (µm) between spatial network neighbors of pairs of cell types aggregated across all edges included in the final set of samples.

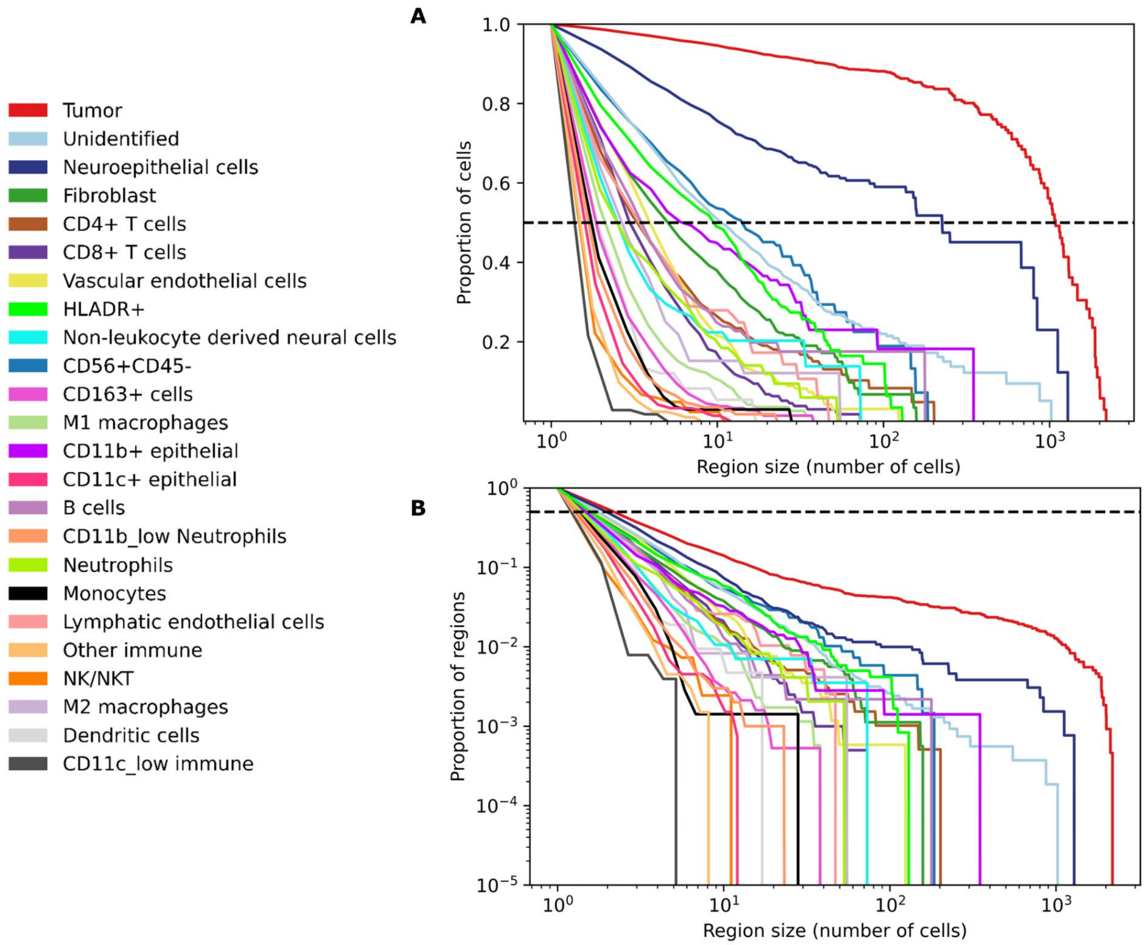


**Fig S4. Cell type region size distributions.** (A) The proportion of cells of each type found within a region of equal or larger size, displayed with a logarithmic x axis. (B) The log-log complementary cumulative distribution function of cell type region sizes aggregated across all connected regions in all samples included in the final analysis.

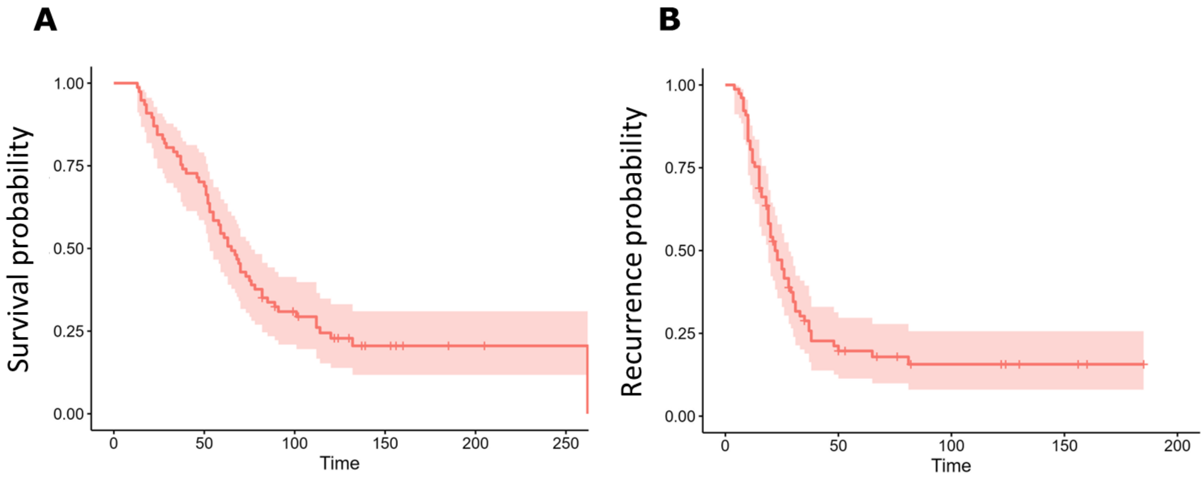


**Fig S5. Survival curves.** Kaplan-Meier survival curves for (A) survival and (B) recurrence outcomes.

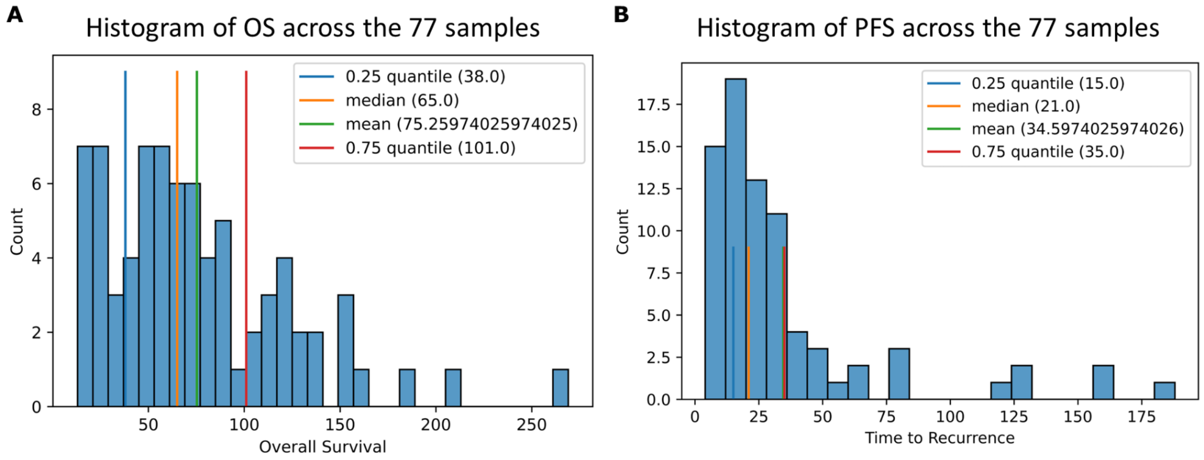


**Fig S6. Outcome variable distributions.** Histograms of the two outcome variables used in our analysis (A) OS, in months, and (B) PFS, in months. For predictive modeling, both outcome variables are split at their respective medians into high and low categories to create a binary prediction task.

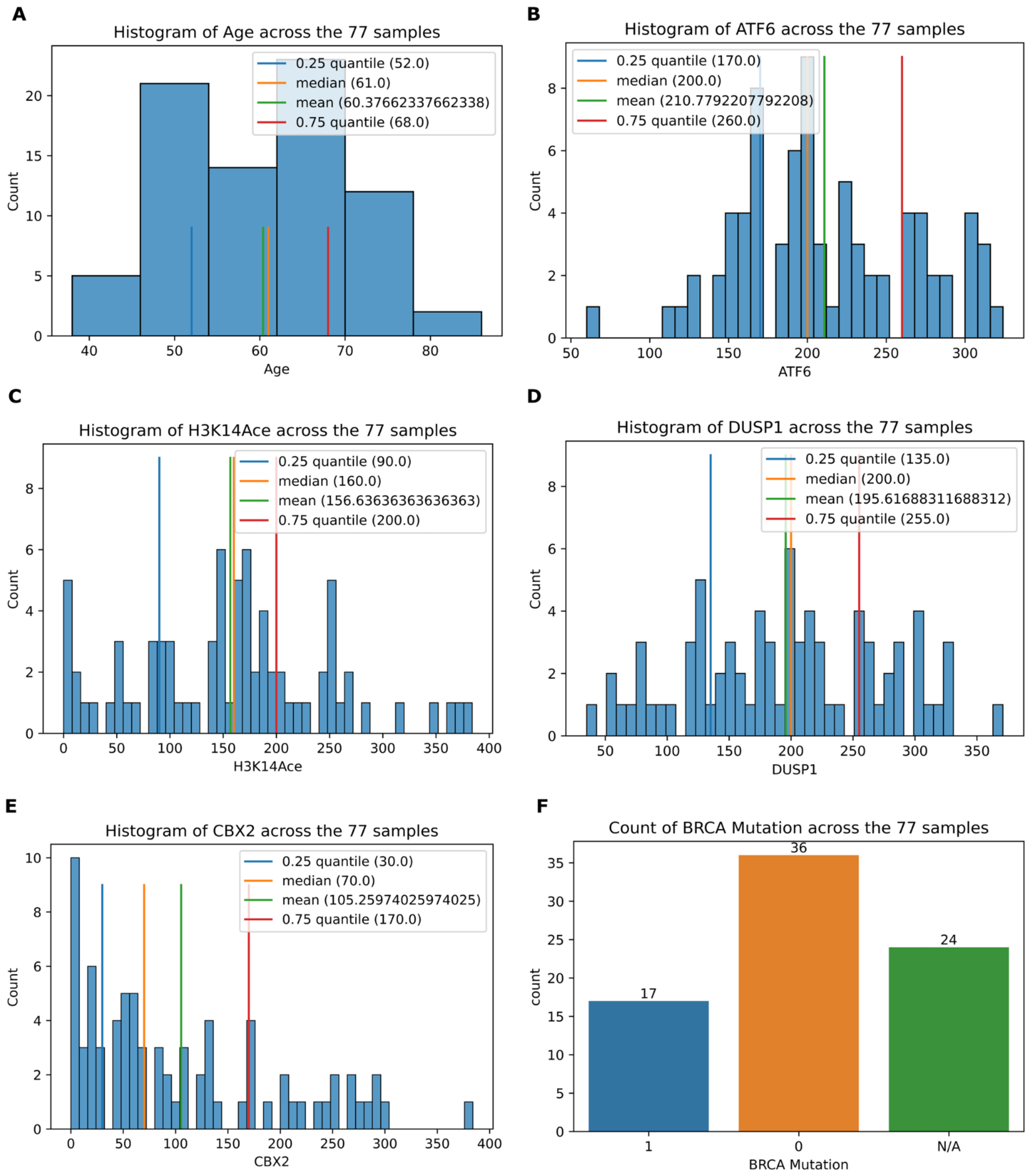


**Fig S7. Clinical/immunohistochemical feature distributions.** Histograms of clinical/immunohistochemical features: (A) age, in years, (B) ATF6 status (histology score), (C) H3K14Ace status (histology score), (D) DUSP1 status (histology score), (E) CBX2 status (histology score), and (F) counts of BRCA Mutation status, where an N/A value indicates that the patient was not tested.

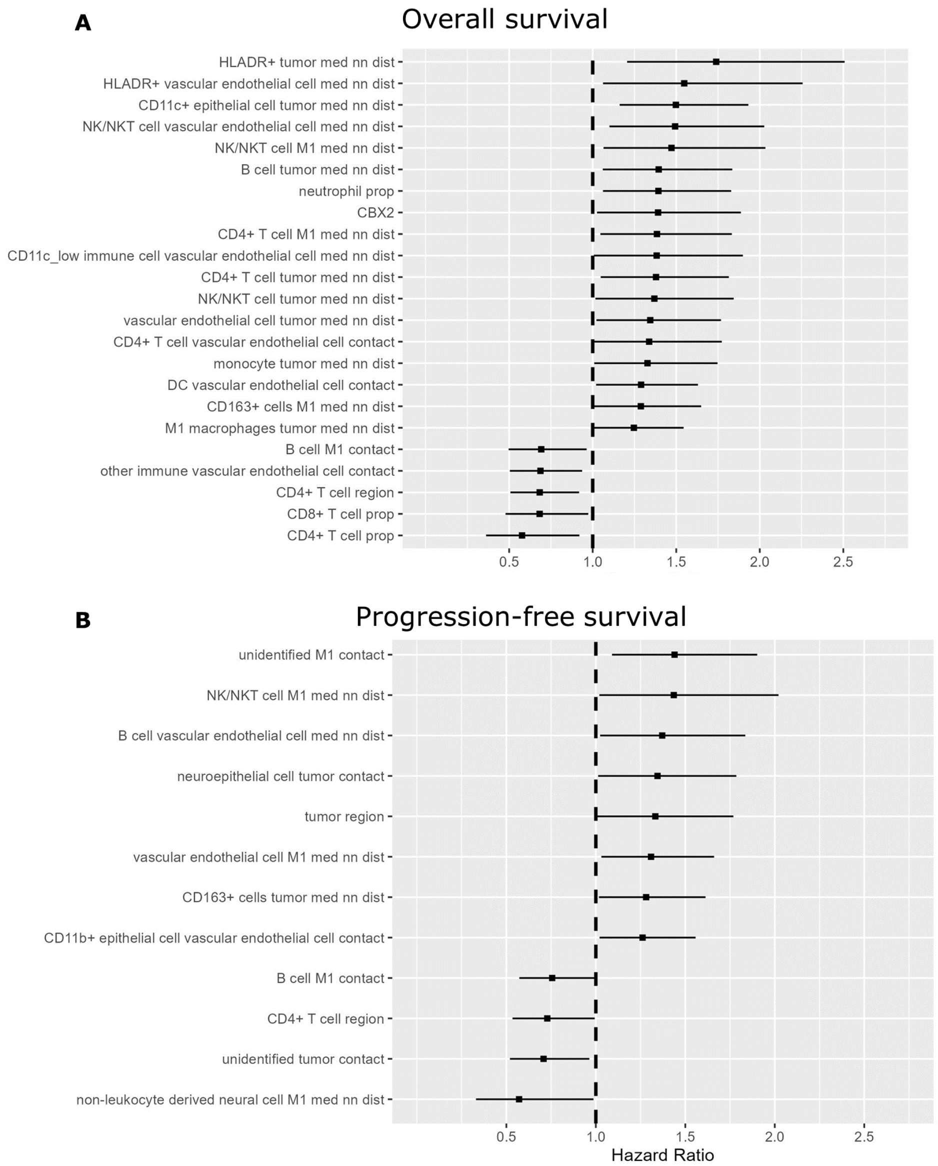


**Fig S8. Univariate Cox regression results for only primary tumor samples.** Covariates found to be significant in Univariate Cox regressions for (A) OS and (B) PFS outcomes. Results are shown for main text Figure 5 if analysis restricted to only primary tumor samples (n=69).

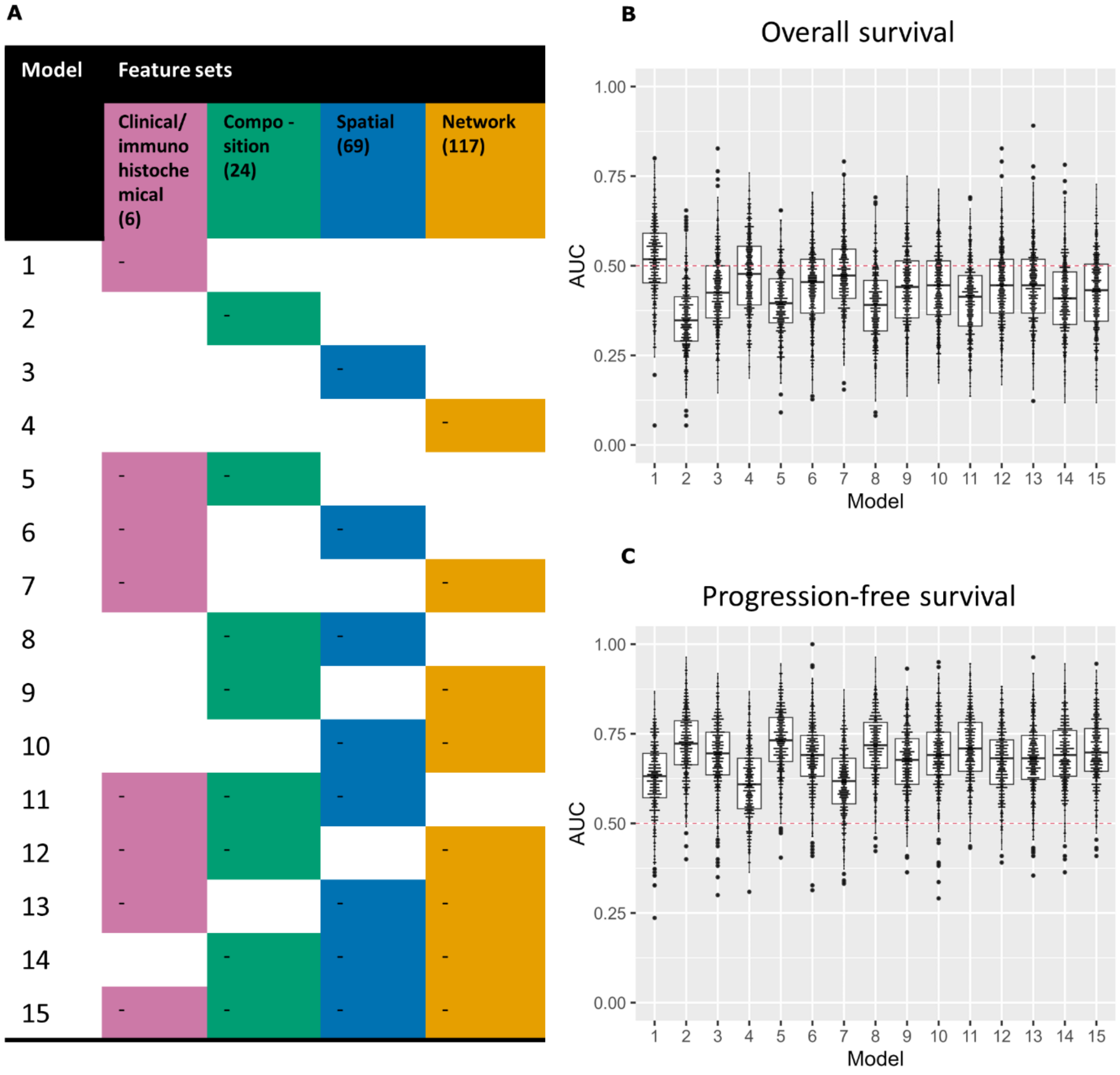


**Fig S9. Radom forest predictive performance results for only primary tumor samples.** Main text Figure 6 repeated with random forest models trained only on primary tumor samples for the same binary outcome variables of low vs. high OS and PFS (n=69).

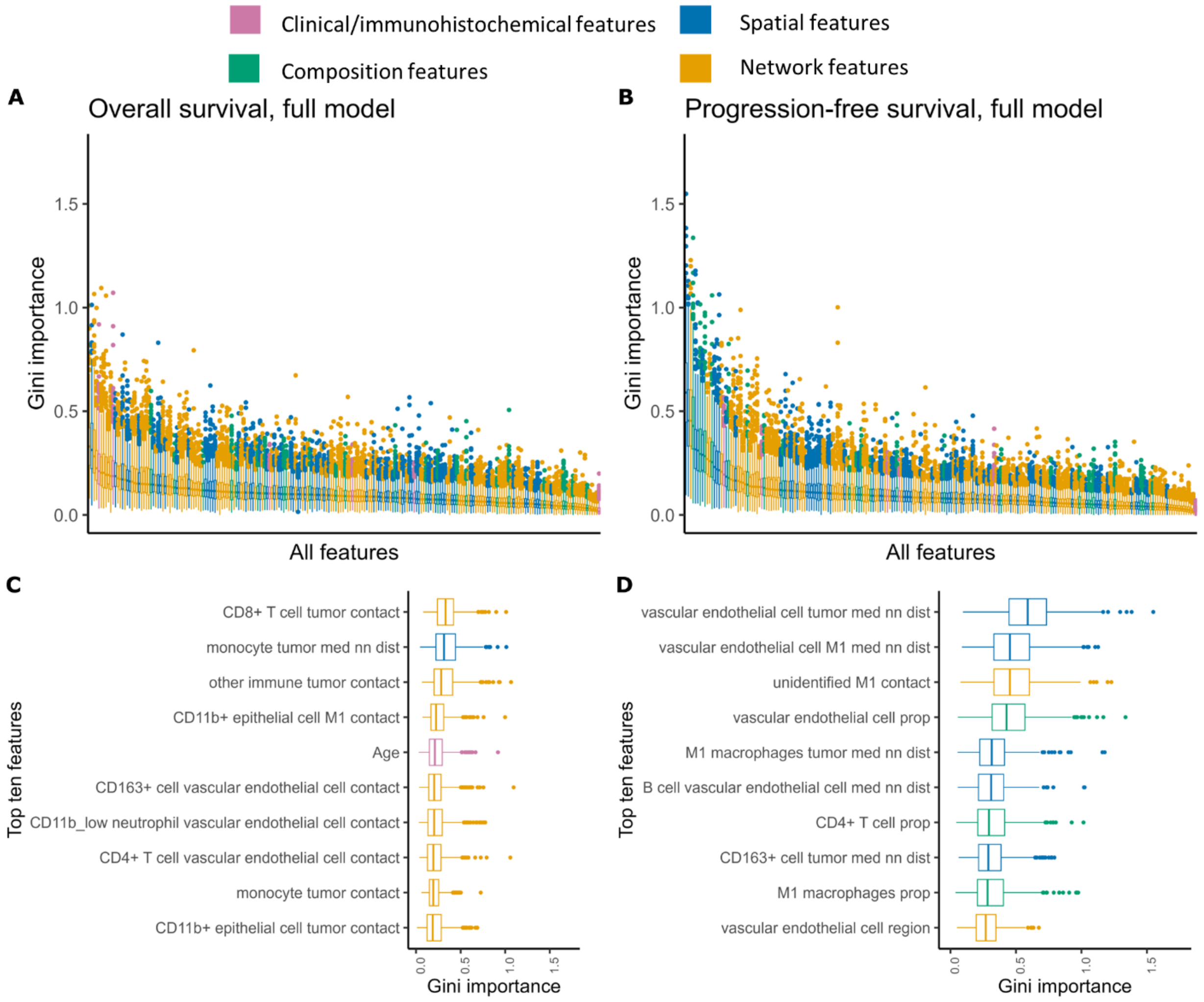


**Fig S10. Random forest feature importance results for only primary tumor samples.** Main text Figure 7 with random forest models trained only on primary tumor samples for the same binary outcome variables of low vs. high OS and PFS (n=69).

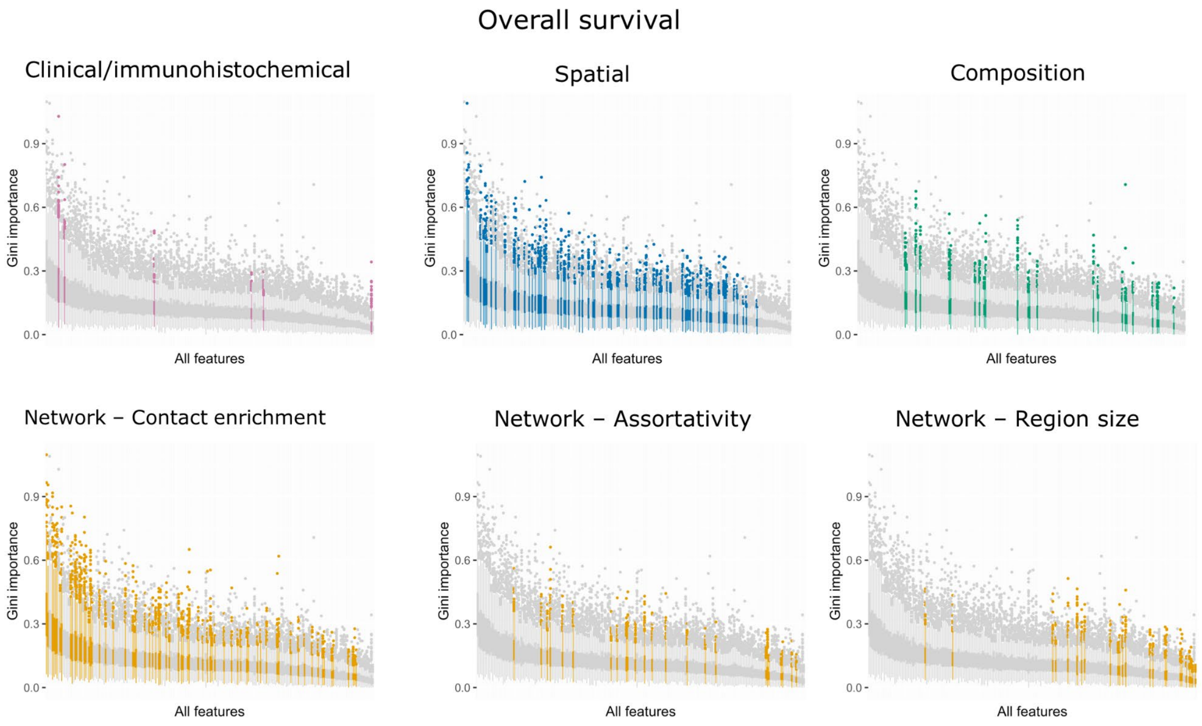


**Fig S11. Random forest feature importance results split by feature type - OS.** Feature importance results for overall survival show in Figure 7A, split by feature type and split by subtype for spatial network features.

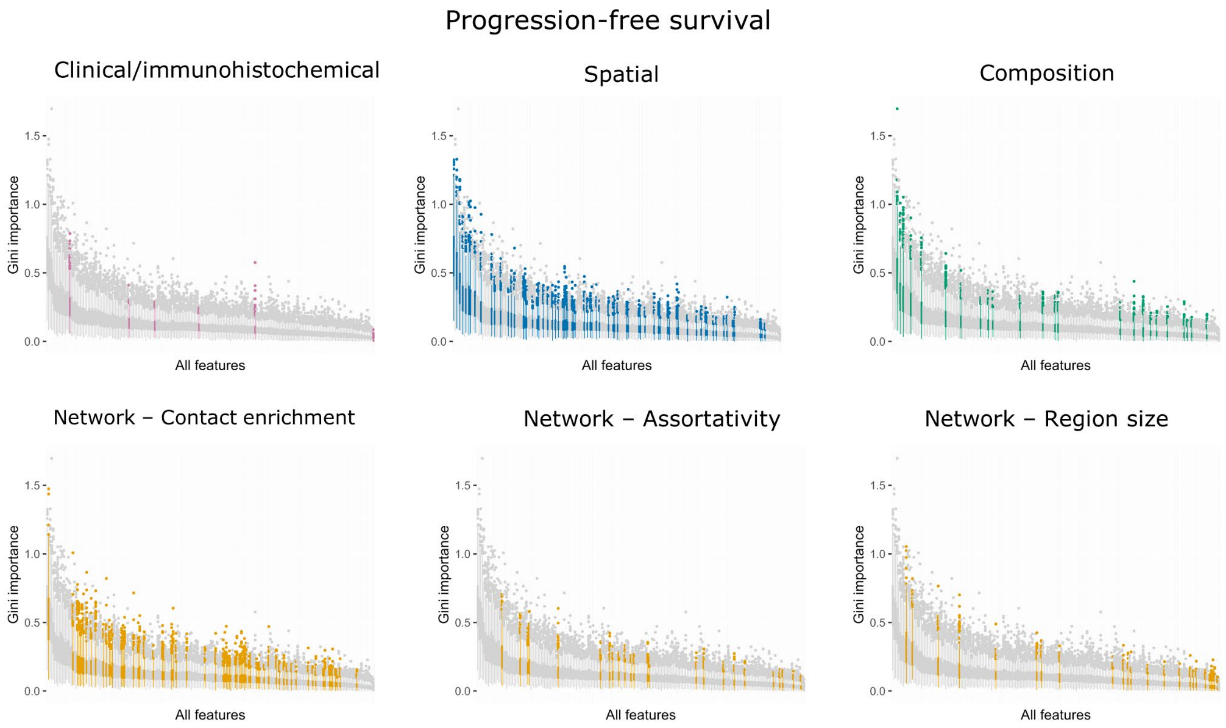


**Fig S12. Random forest feature importance results split by feature type - PFS.** Feature importance results for progression-free survival show in Figure 7B, split by feature type and split by subtype for spatial network features.

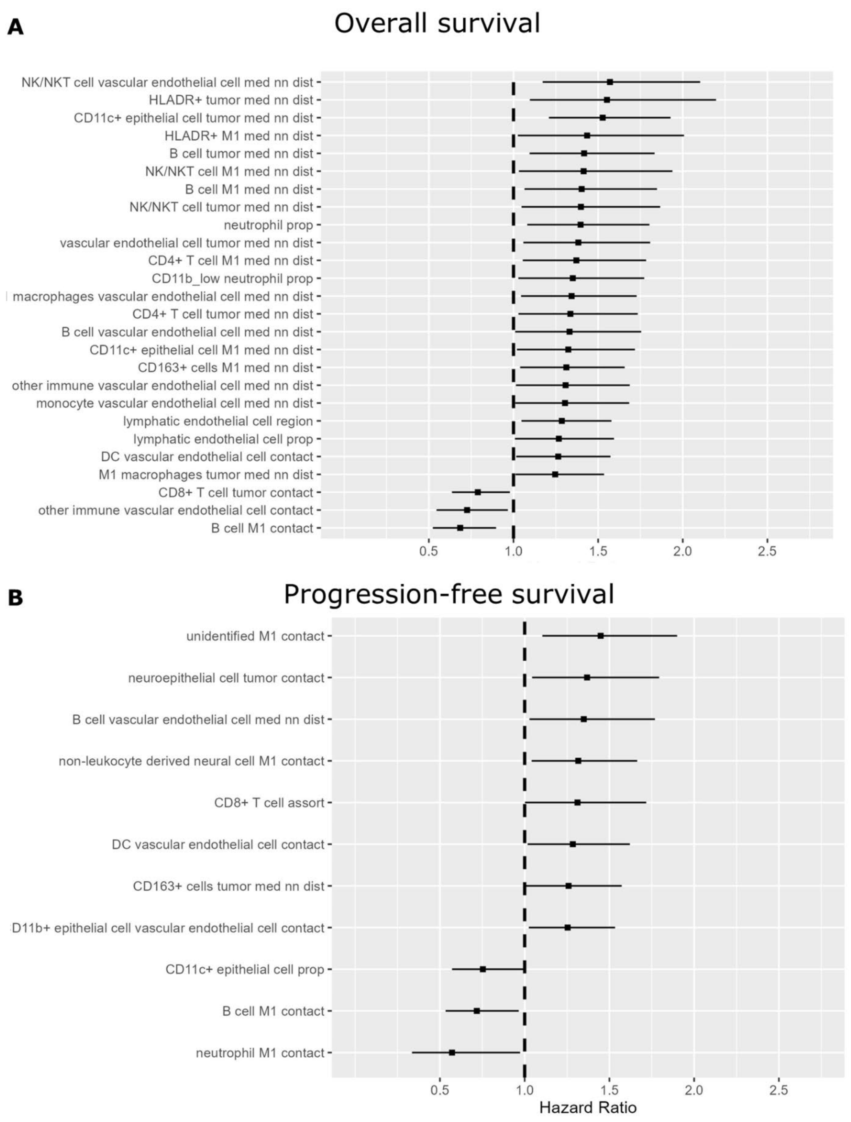


**Fig S13. Univariate Cox regression results with increased spatial network trimming threshold.** Covariates found to be significant in Univariate Cox regressions for (A) OS and (B) PFS outcomes. Results are shown for main text Figure 5 if edges in spatial networks are trimmed with a higher threshold of 100 pixels (~48.8 μm).

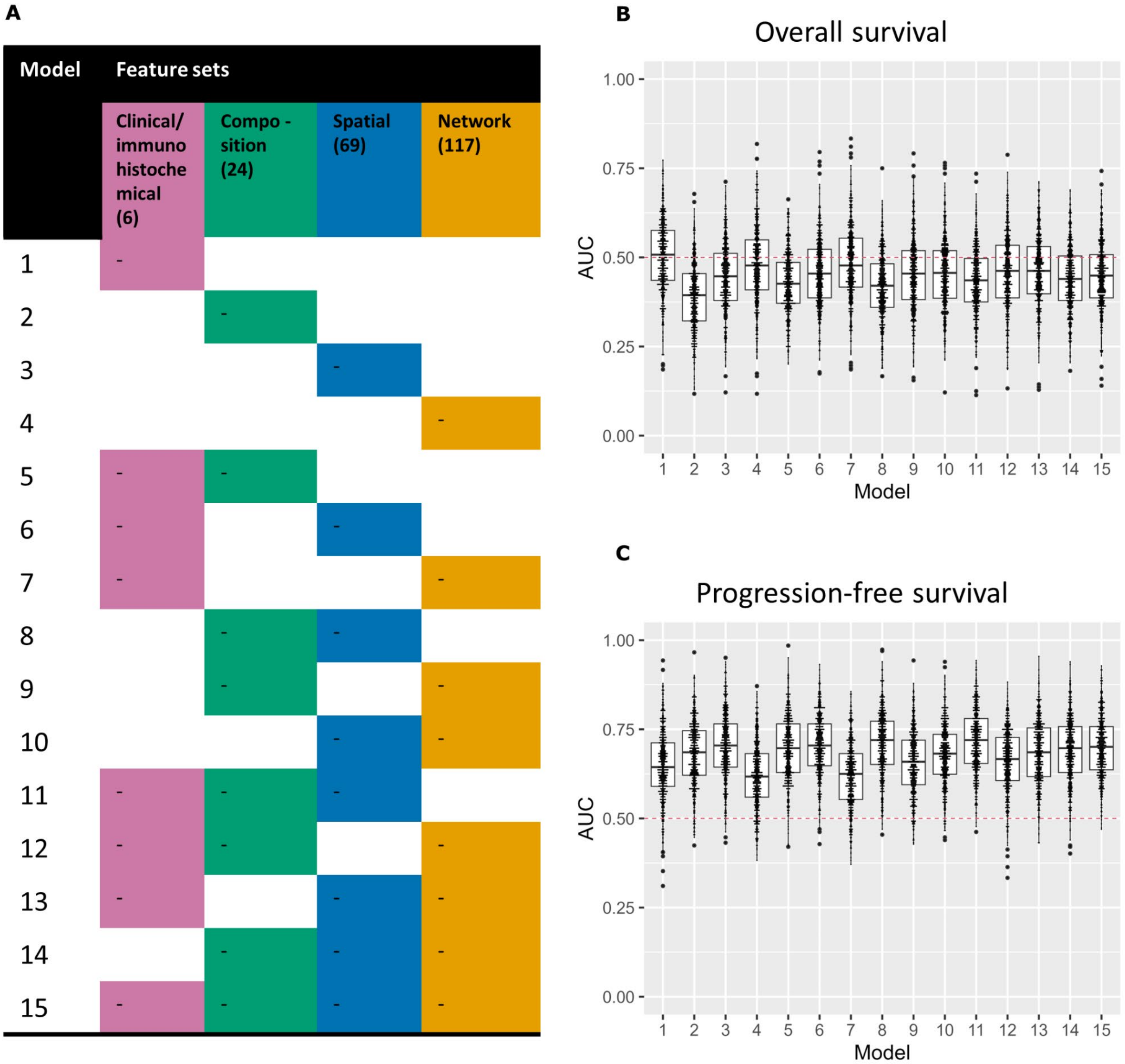


**Fig S14. Random forest predictive performance results with increased spatial network trimming threshold.** Main text Figure 6 repeated if edges in spatial networks are trimmed with a higher threshold of 100 pixels (~48.8 μm).

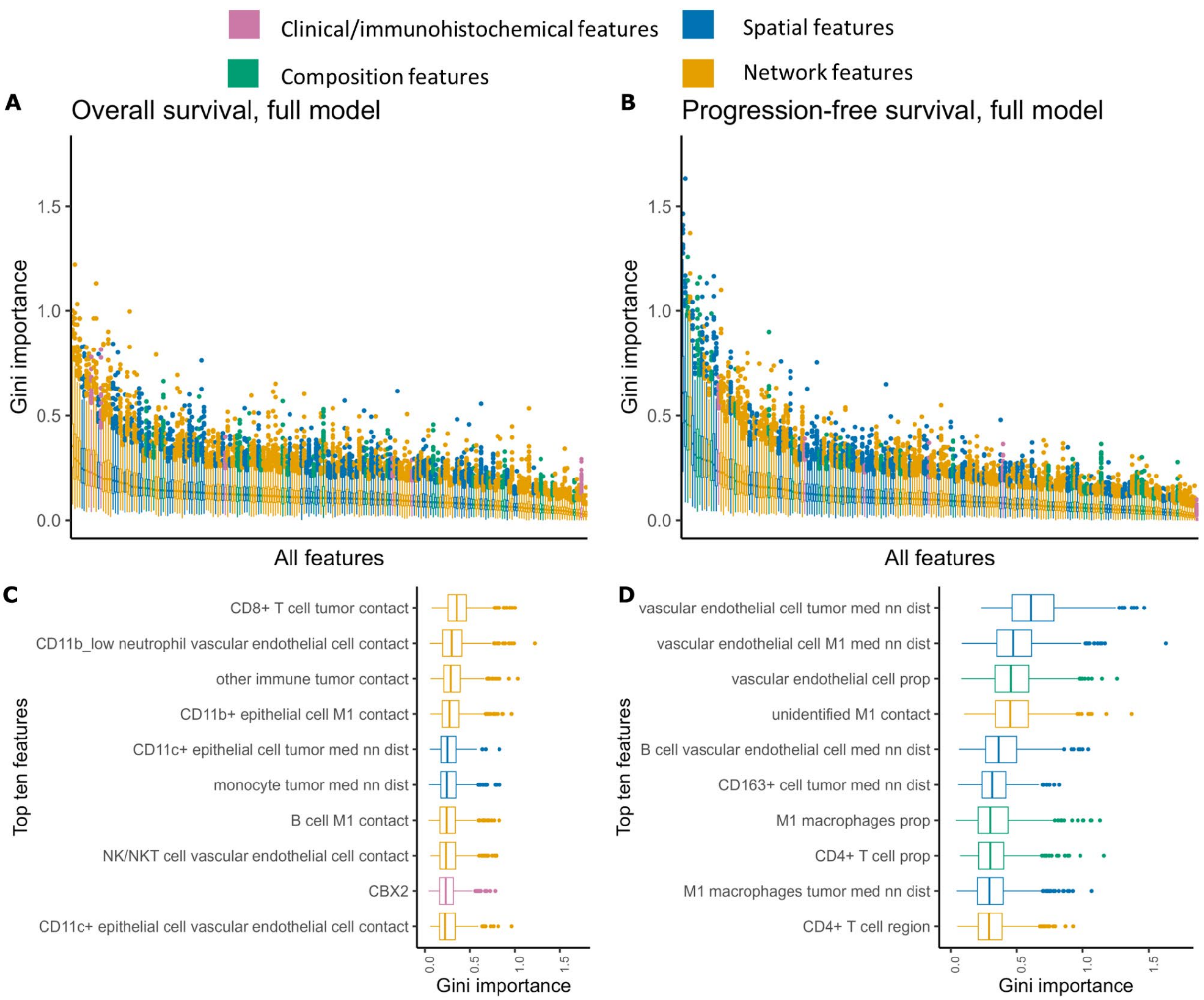


**Fig S15. Random forest feature importance results with increased spatial network trimming threshold.** Main text Figure 7 repeated if edges in spatial networks are trimmed with a higher threshold of 100 pixels (~48.8 μm).

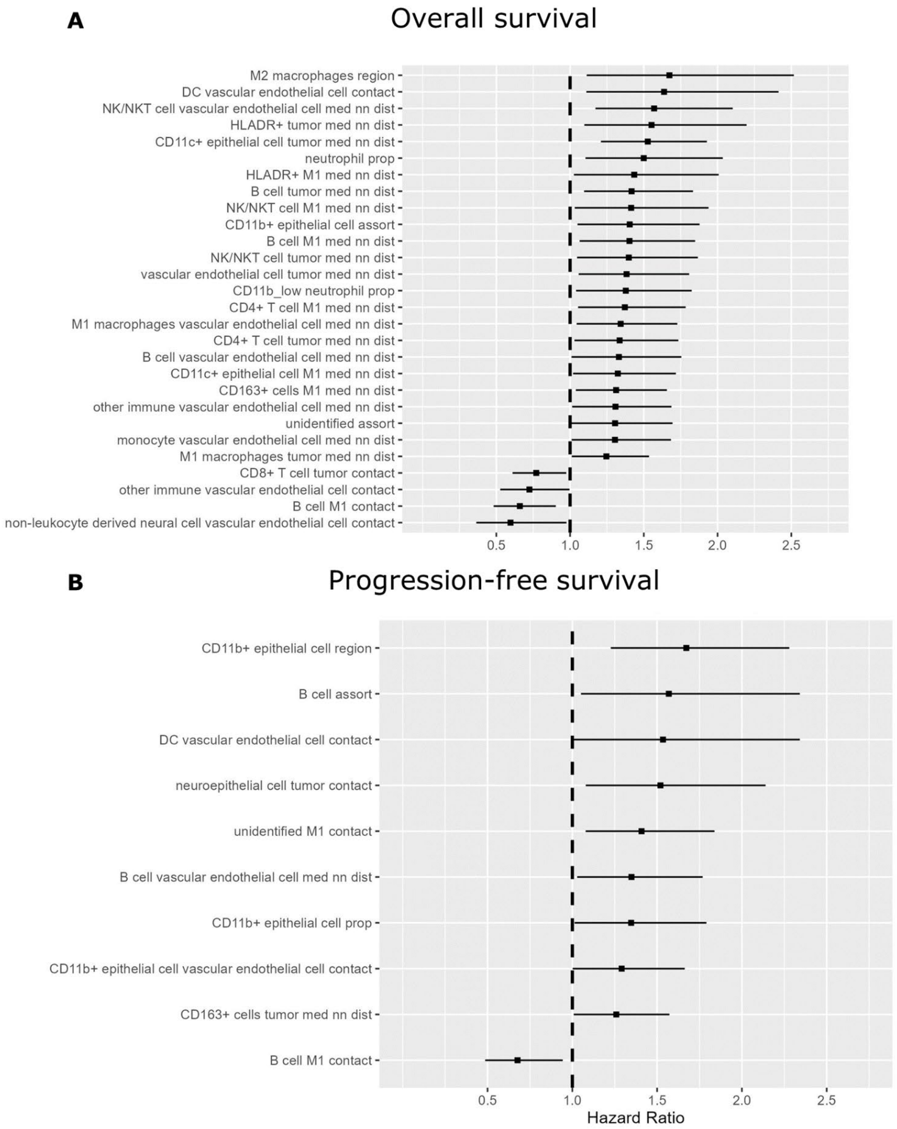


**Fig S16. Univariate Cox regression results with features derived from missing cell types all treated as NA.** Covariates found to be significant in Univariate Cox regressions for (A) OS and (B) PFS outcomes. Results are shown for main text Figure 5 if missing cell types in samples are handled differently in the data pre-processing (see Note S1).

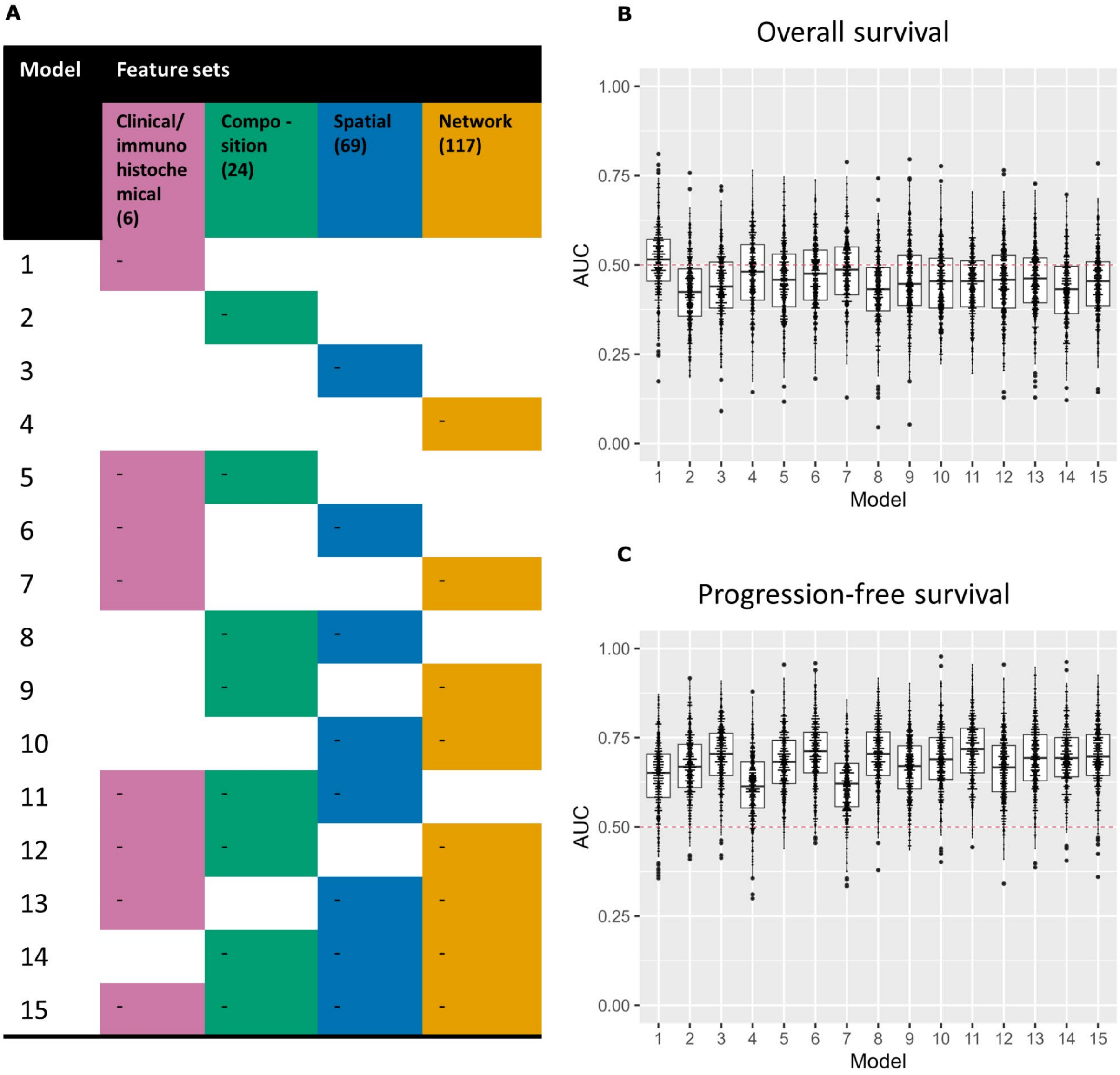


**Fig S17. Random forest predictive performance results with features derived from missing cell types all treated as NA.** Main text Figure 6 repeated with missing cell types handled differently in data preprocessing per Note S1.

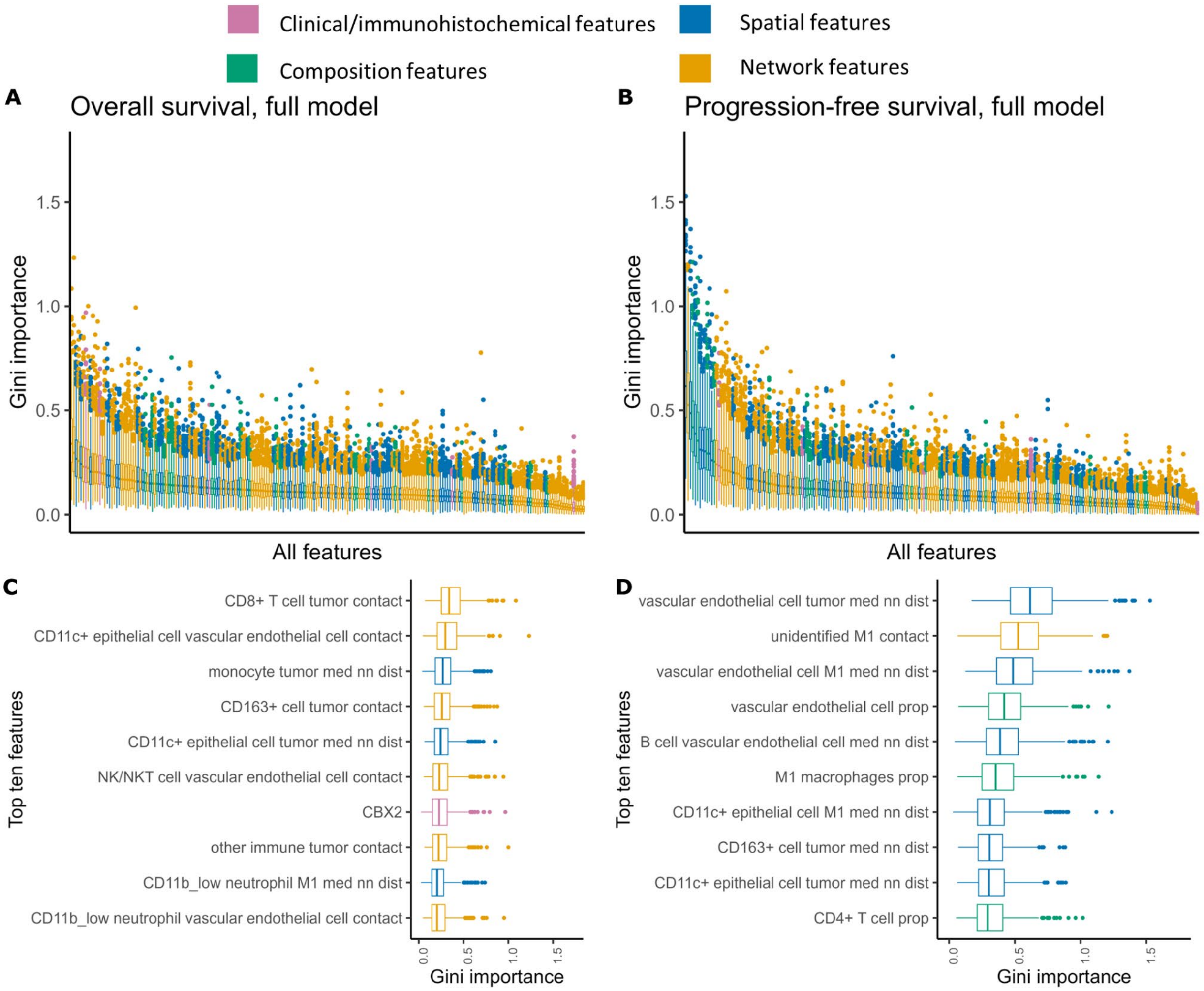


**Fig S18. Random forest feature importance results with features derived from missing cell types all treated as NA.** Main text Figure 7 repeated with missing cell types handled differently in data preprocessing per Note S1.

**Supplementary Tables:**

**Table S1. Mass correction parameters.** Configuration parameters used for multiplexed imaging correction.

| **Recipient Channel** | **Donor Channel** | **Percentage corrected** |
|---|---|---|
| 159 | 142 | 6.9 |
| 158 | 142 | 6.13 |
| 144 | 142 | 0.32 |
| 158 | 156 | 0.96 |
| 175 | 149 | 15 |
| 113 | 89 | 3.3 |
| 142 | 143 | 0.79 |
| 144 | 143 | 2 |
| 149 | 148 | 2.3 |
| 154 | 153 | 3.2 |
| 151 | 153 | 0.8 |
| 165 | 153 | 1 |
| 173 | 172 | 5.2 |
| 173 | 156 | 5 |

**Table S2. Multivariate Cox regression results - OS.** Full multivariate Cox regression results for overall survival, using the top 5 features from univariate analyses. A model controlling for clinical/immunohistochemical features (multivariable) is compared to a model not controlling for these features (multivariable reduced).

| | | **n (%)** | **HR (univariable)** | **HR (multivariable)** | **HR (multivariable reduced)** |
|---|---|---|---|---|---|
| BRCA_Mutation | Negative | 36 (67.9) | - | - | - |
| | Positive | 17 (32.1) | 1.44 (0.75-2.78, p=0.273) | 2.10 (0.56-7.86, p=0.272) | - |
| Age | Mean (SD) | 60.4 (10.3) | 1.02 (0.99-1.04, p=0.207) | 1.01 (0.95-1.07, p=0.861) | - |
| H3K14Ace | Mean (SD) | 156.6 (92.2) | 1.00 (1.00-1.00, p=0.436) | 1.00 (0.99-1.01, p=0.438) | - |
| ATF6 | Mean (SD) | 210.8 (55.6) | 1.00 (1.00-1.01, p=0.333) | 1.00 (0.99-1.01, p=0.980) | - |
| DUSP1 | Mean (SD) | 195.6 (78.0) | 1.00 (1.00-1.00, p=0.857) | 0.99 (0.98-1.00, p=0.075) | - |
| CBX2 | Mean (SD) | 105.3 (95.7) | 1.00 (1.00-1.01, p=0.060) | 1.01 (1.00-1.01, p=0.023) | - |
| CD11c+_epithelial_tumor_med_dist | Mean (SD) | -0.0 (1.0) | 1.53 (1.21-1.93, p<0.001) | 0.46 (0.08-2.62, p=0.385) | 1.38 (0.93-2.06, p=0.110) |
| CD3T_Blood_vessel_med_dist | Mean (SD) | -0.0 (1.0) | 1.57 (1.17-2.10, p=0.002) | 1.71 (0.98-3.00, p=0.061) | 1.66 (1.13-2.42, p=0.009) |
| Lymphatic_vessel_region | Mean (SD) | -0.0 (1.0) | 1.37 (1.09-1.72, p=0.007) | 2.01 (0.98-4.10, p=0.055) | 1.55 (0.94-2.53, p=0.083) |
| B_cells_tumor_med_dist | Mean (SD) | 0.0 (1.0) | 1.42 (1.10-1.83, p=0.008) | 1.67 (0.64-4.36, p=0.294) | 1.29 (0.84-1.98, p=0.251) |
| Neutrophils_prop | Mean (SD) | -0.0 (1.0) | 1.40 (1.08-1.80, p=0.011) | 0.02 (0.00-7.76, p=0.194) | 1.11 (0.78-1.59, p=0.558) |

**Table S3. Multivariate Cox regression results - PFS.** Full multivariate Cox regression results for progression-free survival, using the top 5 features from univariate analyses. A model controlling for clinical/immunohistochemical features (multivariable) is compared to a model not controlling for these features (multivariable reduced).

| | | **n (%)** | **HR (univariable)** | **HR (multivariable)** | **HR (multivariable reduced)** |
|---|---|---|---|---|---|
| BRCA_Mutation | Negative | 36 (67.9) | - | - | - |
| | Positive | 17 (32.1) | 0.99 (0.52-1.86, p=0.966) | 0.67 (0.26-1.75, p=0.417) | - |
| Age | Mean (SD) | 60.4 (10.3) | 1.00 (0.98-1.03, p=0.812) | 1.01 (0.97-1.05, p=0.628) | - |
| H3K14Ace | Mean (SD) | 156.6 (92.2) | 1.00 (1.00-1.00, p=0.247) | 1.00 (1.00-1.01, p=0.444) | - |
| ATF6 | Mean (SD) | 210.8 (55.6) | 1.00 (1.00-1.01, p=0.201) | 1.00 (1.00-1.01, p=0.376) | - |
| DUSP1 | Mean (SD) | 195.6 (78.0) | 1.00 (1.00-1.00, p=0.333) | 0.99 (0.99-1.00, p=0.085) | - |
| CBX2 | Mean (SD) | 105.3 (95.7) | 1.00 (1.00-1.00, p=0.281) | 1.00 (0.99-1.00, p=0.828) | - |
| Unidentified_M1_contact | Mean (SD) | -0.0 (1.0) | 1.41 (1.08-1.84, p=0.010) | 1.63 (1.12-2.36, p=0.010) | 1.32 (0.99-1.76, p=0.059) |
| Neuroepithelial_cells_tumor_contact | Mean (SD) | 0.0 (1.0) | 1.40 (1.07-1.83, p=0.015) | 1.64 (0.96-2.80, p=0.071) | 1.40 (0.98-1.99, p=0.066) |
| CD11b+_epithelial_Blood_vessel_contact | Mean (SD) | -0.0 (1.0) | 1.25 (1.02-1.53, p=0.029) | 1.25 (0.88-1.76, p=0.209) | 1.31 (1.02-1.66, p=0.031) |
| B_cells_Blood_vessel_med_dist | Mean (SD) | -0.0 (1.0) | 1.35 (1.03-1.77, p=0.030) | 1.40 (0.92-2.11, p=0.114) | 1.56 (1.15-2.12, p=0.004) |

| | | | | | |
|---|---|---|---|---|---|
| CD8T_assort | Mean (SD) | 0.0 (1.0) | 1.34 (1.02-1.76, p=0.034) | 1.50 (0.96-2.33, p=0.072) | 1.70 (1.20-2.41, p=0.003) |

**Table S4. Random forest predictive performance results.** Full results for all random forest models using 15 different feature sets. Mean and standard deviation ROC-AUC results across 500 evaluations are reported for both OS and PFS.

| model | Feature set size | mean.OS | sd.OS | mean.PFS | sd.PFS |
|---|---|---|---|---|---|
| 1 | 6 | 0.501772727 | 0.097896911 | 0.645575758 | 0.098999496 |
| 2 | 24 | 0.394969697 | 0.088970605 | 0.677106061 | 0.087990041 |
| 3 | 69 | 0.437356061 | 0.094588399 | 0.703310606 | 0.082421898 |
| 4 | 117 | 0.438007576 | 0.104566875 | 0.616984848 | 0.090243279 |
| 5 | 30 | 0.422371212 | 0.095970962 | 0.697613636 | 0.09284265 |
| 6 | 75 | 0.454318182 | 0.093963455 | 0.697090909 | 0.086857852 |
| 7 | 123 | 0.453189394 | 0.104417303 | 0.625681818 | 0.096502911 |
| 8 | 93 | 0.410606061 | 0.095374843 | 0.710939394 | 0.087193185 |
| 9 | 141 | 0.426871212 | 0.107178202 | 0.662886364 | 0.085802621 |
| 10 | 186 | 0.430590909 | 0.094591019 | 0.685712121 | 0.085374049 |
| 11 | 99 | 0.432075758 | 0.095595225 | 0.706833333 | 0.088637524 |
| 12 | 147 | 0.429977273 | 0.102141897 | 0.669734848 | 0.087931972 |
| 13 | 192 | 0.441848485 | 0.095468565 | 0.687901515 | 0.091137068 |
| 14 | 210 | 0.424772727 | 0.098449836 | 0.696568182 | 0.090182419 |
| 15 | 216 | 0.434469697 | 0.096037455 | 0.696583333 | 0.08819006 |

**Table S5. Random forest predictive performance results for only primary tumor samples.** Full results for all random forest models using 15 different feature sets, trained on only primary tumor samples (n=69). Mean and standard deviation ROC-AUC results across 500 evaluations are reported for both OS and PFS.

| model | feature_set_sizes | mean.Survival | sd.Survival | mean.Recurrence | sd.Recurrence |
|---|---|---|---|---|---|
| 1 | 6 | 0.516882 | 0.108072 | 0.625582 | 0.099834 |
| 2 | 24 | 0.351182 | 0.096781 | 0.719336 | 0.093301 |
| 3 | 69 | 0.427273 | 0.10353 | 0.688473 | 0.096683 |
| 4 | 117 | 0.474355 | 0.109261 | 0.613036 | 0.099609 |
| 5 | 30 | 0.398309 | 0.093621 | 0.728718 | 0.088445 |
| 6 | 75 | 0.444482 | 0.109681 | 0.688782 | 0.094147 |
| 7 | 123 | 0.479136 | 0.105266 | 0.616482 | 0.097207 |
| 8 | 93 | 0.389682 | 0.101541 | 0.714309 | 0.091051 |
| 9 | 141 | 0.437964 | 0.11196 | 0.672536 | 0.08995 |
| 10 | 186 | 0.439027 | 0.106971 | 0.6925 | 0.094976 |
| 11 | 99 | 0.405282 | 0.104064 | 0.711936 | 0.093711 |
| 12 | 147 | 0.446873 | 0.109477 | 0.673191 | 0.090713 |
| 13 | 192 | 0.445145 | 0.110672 | 0.680245 | 0.096609 |
| 14 | 210 | 0.410682 | 0.106634 | 0.691236 | 0.090375 |
| 15 | 216 | 0.426464 | 0.108707 | 0.700445 | 0.092391 |

**Table S6. Definitions of cellular phenotypes identified with unsupervised clustering**.

| Phenotype | Differential Markers[1] | Lineage |
|---|---|---|
| CD4+ T cells | CD3+, CD4+, CD45+ | Lymphocyte |
| CD8+ T cells | CD3+, CD8+, CD45+ | Lymphocyte |
| B cells | CD20+, CD45+ | Lymphocyte |
| NK/NKT | CD3+/-, CD56+, CD45+, | NK = CD56+/CD45+/CD3- |

| | | |
|---|---|---|
| | | NKT = CD56+/CD45+/CD3+ Lymphocyte |
| M2 macrophages | CD68+, CD163+, CD45+ | Myeloid |
| M1 macrophages | CD68+, CD163-, CD45+ | Myeloid |
| Dendritic cells | DC-SIGN+, CD45+ | Myeloid |
| Monocyte | CD45+, CD11b+, CD56- | Myeloid |
| $CD11c^{low}$ immune | CD45+, $CD11c^{low}$ | Myeloid |
| Other immune | CD45+ | Leukocytes |
| Neutrophils | CD45-, CD11b+ | Granulocyte |
| $CD11b^{low}$ Neutrophils | CD45-, $CD11b^{low}$, $CD11c^{low}$ | Granulocyte |
| Vascular endothelial cells | CD31+, CD45- | Endothelial |
| Lymphatic endothelial cells | Podoplanin+, CD45- | Endothelial |
| Fibroblast | Vimentin+, CD45- | Mesenchymal |
| Non-leukocyte derived neural cells | CD56+, CD45-, Keratin- | Neural cell |
| CD56+CD45- | CD56+, CD45-, Vimentin- | Neural cell |
| Neuroepithelial cells | CD56+, Keratin+, CD45- | Epithelial |
| CD11b+ epithelial | CD11b+, Keratin+, CD45- | Epithelial |
| CD11c+ epithelial | $CD11c^{low}$, $CD11b^{low}$, Keratin+ | Epithelial |
| Tumor | Keratin+ | Epithelial |
| HLADR+ | HLADR+, CD45- | No expression of other phenotypic markers |
| CD163+ | CD45+/-, CD163+, CD11b+/-, CD11c+/- | Mixed |

| | | |
|---|---|---|
| Unidentified | -/- | No expression of phenotypic markers |
| [1]Markers that differentiate population clusters are listed. Refer to the heatmap in Figure 1C for the description of all phenotypic markers. | | |

### References

1. Newman MEJ. Mixing patterns in networks. Phys Rev E. 2003;67:026126.

2. Forman G, Scholz M. Apples-to-apples in cross-validation studies: pitfalls in classifier performance measurement. SIGKDD Explor Newsl. 2010;12:49–57.
